# Allosteric control of Ubp6 and the proteasome via a bidirectional switch

**DOI:** 10.1101/2021.07.16.452743

**Authors:** Ka Ying Sharon Hung, Sven Klumpe, Markus R. Eisele, Suzanne Elsasser, Geng Tian, Shuangwu Sun, Jamie A. Moroco, Tat Cheung Cheng, Tapan Joshi, Timo Seibel, Duco Van Dalen, Xin-Hua Feng, Ying Lu, Huib Ovaa, John R. Engen, Byung-Hoon Lee, Till Rudack, Eri Sakata, Daniel Finley

**Affiliations:** Department of Cell Biology, Harvard Medical School, Boston, MA 02115, USA; Department of Molecular Structural Biology, Max Planck Institute of Biochemistry, 82152 Martinsried, Germany; Department of Biochemistry, Gene Center, Ludwig-Maximilians-Universität, Munich, 81377 Munich, Germany; Life Sciences Institute (LSI), Zhejiang University, Hangzhou, China 310058; Department of Chemistry and Chemical Biology, Northeastern University, Boston, MA 02115, USA; Institute for Auditory Neuroscience, University Medical Center Göttingen, 37077 Göttingen, Germany; Leiden University Medical Center, Einthovenweg 20, 2333 ZC Leiden the Netherlands; Department of Systems Biology, Harvard Medical School, Boston, MA 02115, USA; Department of New Biology, Daegu Gyeongbuk Institute of Science and Technology (DGIST), Daegu 42988, Korea; Biospectroscopy, Center for Protein Diagnostics (PRODI), Ruhr University Bochum, 44801 Bochum, Germany; Department of Biophysics, Ruhr University Bochum, 44801 Bochum, Germany

## Abstract

The proteasome is the principal cellular protease, and recognizes target proteins that have been covalently marked by ubiquitin chains. The ubiquitin signal is subject to rapid editing at the proteasome, allowing it to reject substrates based on topological features of their attached ubiquitin chains. Editing is mediated by a key regulator of the proteasome, deubiquitinating enzyme Ubp6. The proteasome activates Ubp6, whereas Ubp6 inhibits the proteasome–both by deubiquitinating proteasome-bound ubiquitin conjugates, and through a noncatalytic effect that does not involve deubiquitination. We report mutants in both Ubp6 and proteasome subunit Rpt1 that abrogate Ubp6 activation. The Ubp6 mutations fall within its ILR element, defined here, which is conserved from yeast to mammals. The ILR is a component of the BL1 blocking loop, other parts of which obstruct ubiquitin access to the catalytic groove in free Ubp6. Rpt1 docking at the ILR opens the catalytic groove by rearranging not only BL1 but also a novel network of three directly interconnected active-site-blocking loops. Ubp6 activation and noncatalytic proteasome inhibition by Ubp6 are linked in that they were eliminated by the same Ubp6 and Rpt1 mutations. Ubp6 and ubiquitin together drive the proteasome into a unique conformational state associated with proteasome inhibition. Our results identify a multicomponent allosteric switch that exerts simultaneous control over the activity of both Ubp6 and the proteasome, and suggest that their active states are in general mutually exclusive. The findings lead to a new paradigm for allosteric control of deubiquitinating enzymes.

## Main text

Ubiquitinated substrates are recognized by the 19-subunit regulatory particle (RP) of the proteasome, then translocated by an ATPase motor through a narrow channel into the proteasome core particle (CP) to be degraded. Because ubiquitin modifications constitute a kinetic impediment to translocation^1–6^, substrates are typically deubiquitinated prior to the completion of translocation. Release of ubiquitin both spares it from degradation and provides a checkpoint for the control of proteasome output^7^. Deubiquitinating enzyme Ubp6 (in mammals, USP14) associates with the proteasome reversibly, and suppresses degradation through kinetic competition with the proteasome. With a favorable *in vitro* substrate, ubiquitin removal by Ubp6 is detected in less than a second–before the proteasome initiates degradation^8^. Here we investigate the mechanism by which Ubp6 is activated by the proteasome.

### Ubp6 mutants defective in activation

We performed cryo-EM analysis of the proteasome complexed to Ubp6 covalently bound through its active site cysteine to ubiquitin-vinyl-sulfone (UbVS), achieving substantially greater resolution than previously reported^9^ for this complex (Fig. 1a, Extended Data Fig. 1, Supplementary Table 1). Details of this ternary complex will be presented below; we will focus initially on targeted mutagenesis based on this structure. Ubp6 has two domains: an N-terminal ubiquitin-like (UBL) domain, which binds the proteasome via subunit Rpn1^10–12^, and a C-terminal catalytic domain, which contacts Rpt1 (Extended Data Fig. 2a). The Rpt1 contact site^9^ (Fig. 1b) comprises Interfaces A (R316-V333 of Ubp6 and G158-E169 of Rpt1) and B (E473-S488 of Ubp6 and Y181-R190 of Rpt1). We validated Interface A by showing that it is uniquely protected from hydrogen-deuterium exchange within Ubp6-ubiquitin-vinyl-methyl-ester (UbVME)-RP ternary complexes (Fig. 1c, Extended Data Fig. 2b,c).

**Fig. 1.**
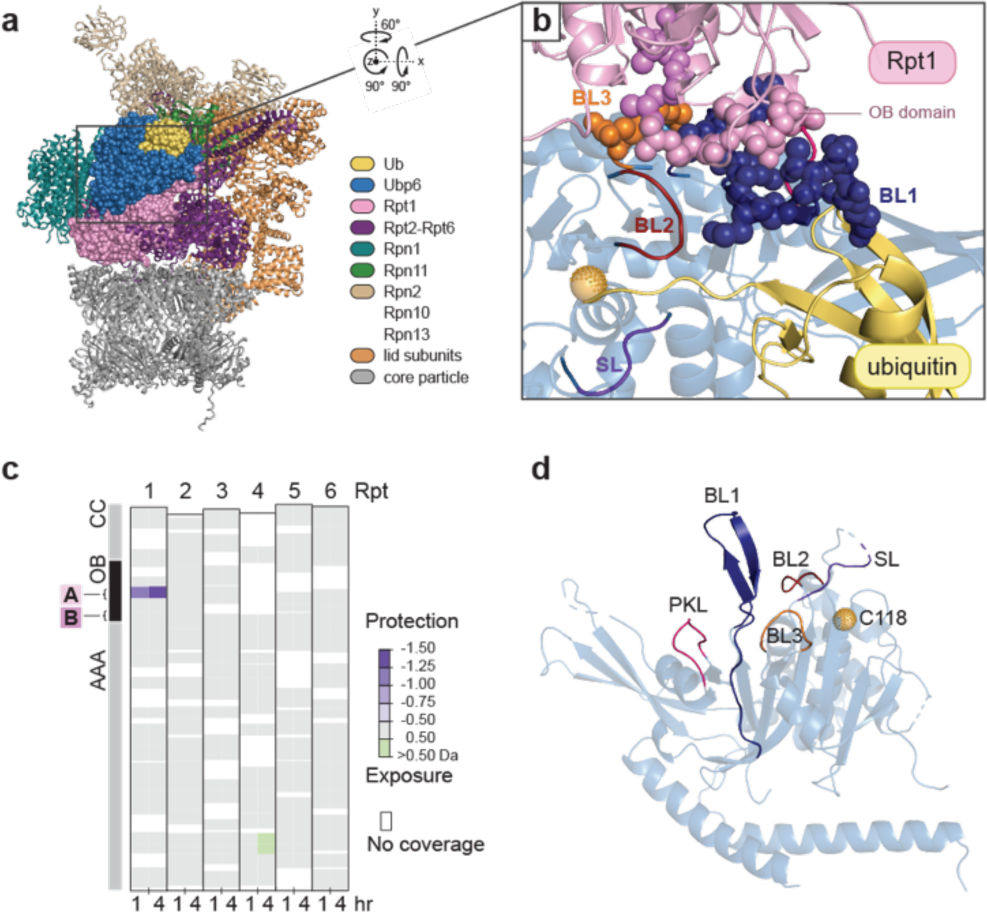
Model of the Ubp6-Rpt1 interface. **a**, Model of the Ubp6-UbVS-proteasome complex based on cryo-EM analysis. Ubp6, Rpt1, and ubiquitin are highlighted as spheres. Boxed region is rotated and enlarged in *b.* (PDB: this study). **b**, The Ubp6-Rpt1 interface with mutagenized residues rendered as spheres. Interface A is in blue for Ubp6 and pink for Rpt1; Interface B, orange for Ubp6, violet for Rpt1. **c**, Interface A is protected from hydrogen exchange by the addition of excess Ubp6-UbVME. Ubp6-dependent deuteration differences within peptides of Rpt1-Rpt6 of purified regulatory particle are shown. See Extended Data Fig. 2 for additional HXMS data. **d**, Critical loops of free Ubp6 (PDB: 1VJV).

**Fig. 2.**
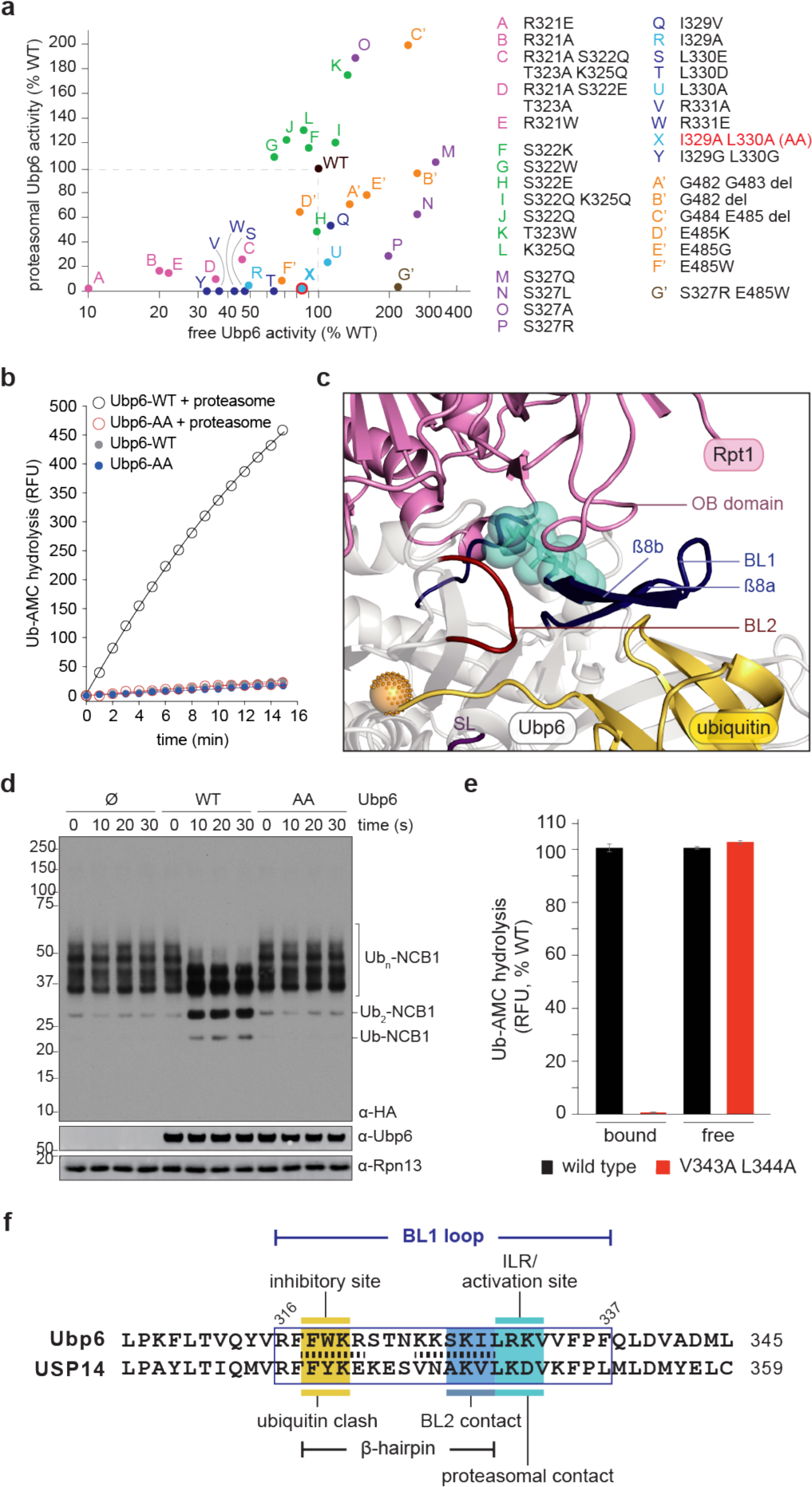
Ubp6 and USP14 mutants that are refractory to proteasome activation. **a**, Activity of Ubp6 mutants in the presence or absence of proteasome, with Ubp6-AA highlighted. **b**, Ub-AMC hydrolysis by Ubp6-AA (60 nM). **c**, Positions of I329 and L330 (cyan) in the modeled Ubp6-Rpt1 interface. (PDB: this study). **d**, Activity of Ubp6-AA on HA-Ubn-NCB1 in the presence of proteasome (and ADP to prevent substrate degradation). No deubiquitination was seen without proteasome (data not shown). **e**, Activity of the USP14-V343A L344A with or without proteasome. Wild-type values are independently set to 100% but differ by over 100-fold, reflecting the extent of activation. Mean ± s.d. is shown. **f**, Sequence alignment between Ubp6 and USP14. Ubiquitin clash for free Ubp6 is shown in Extended Data Fig. 3a. Stippled lines, β-strand-forming residues in Ubp6 (see Extended Data Fig. 3c). Fig. 5e describes BL2 contacts.

Interface A of Ubp6 lies within blocking loop 1 (BL1), which, in the absence of the proteasome, occludes the active site groove of the enzyme^13^ (Extended Data Figs. 2a, 3a). BL1 is rearranged to interpose between ubiquitin and the proteasome in the ternary complex (Fig. 1b). Interface B of Ubp6 is formed by the BL3 loop (Fig. 1b, Extended Data Fig. 2a). BL3 does not occlude the active site of free Ubp6 but rather contacts BL1 at its foot, as does the PKL loop on the opposite side of BL1 (Fig. 1d). BL1 is thus immobilized in free Ubp6. For ubiquitin to access the active site, two additional loops, BL2 and switching loop (SL), must be displaced (Extended Data Fig. 3a). However, unlike BL1 these loops are not in contact with the proteasome (Fig. 1b), suggesting that BL1 may convey the proteasome’s signal for activation to these elements. We refer to BL1, BL2, and SL collectively as the blocking loops. They are all strongly conserved in evolution, indicating their functional importance (Extended Data Fig. 3b).

**Fig. 3.**
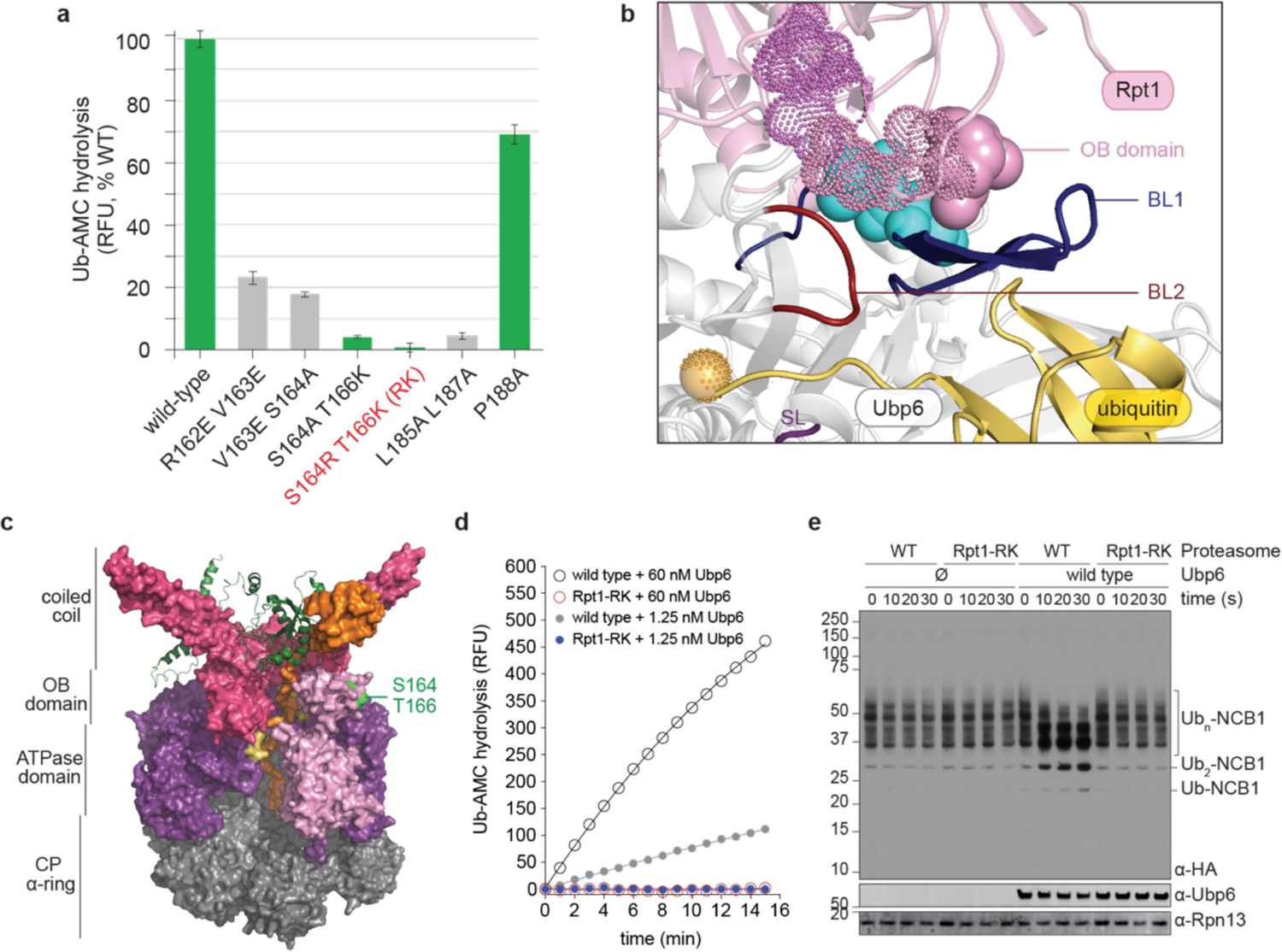
An Rpt1 mutant defective in Ubp6 activation. **a**, Ub-AMC hydrolysis by Ubp6 in the presence of wild-type and mutant proteasomes. Mean ± s.d. is shown. *rpt1-RK* is highlighted. Mutants in green showed unperturbed proteasome assembly (see Extended Data Fig. 5); assembly was impaired for those in grey. **b**, The modeled Rpt1-Ubp6 interface. Cyan, Ubp6 residues I329 and L330. Solid pink, Rpt1 S164 and T166. Stippled pink and violet spheres, other mutated Rpt1 residues (PDB: this study). **c**, Positions of S164 and T166 (neon green) within the OB domain of the ATPase ring. Coiled-coil and OB domains are in rose, ATPase domain in purple, Rpt1 in pink, Rpn11 in green ribbon. Substrate (orange) is passing through RPT pore loops (yellow) into the CP axial channel. Portions of Rpt1 and Rpt2 were removed to reveal the substrate channel. (PDB: 6EF3). **d**, OB domain, top view. Rpt1’s L34 loop is in black, S164 and T166 in green. (PDB: 6EF3). **e**, Ubp6 activity in the presence of mutant proteasomes. **f**, Deubiquitination of HA-Ub_n_-NCB1 in the presence of Ubp6 and ADP-proteasomes.

Interfaces A and B were chosen for mutagenesis (Fig. 2a). We screened for mutants in which the activity of proteasome-bound Ubp6 is reduced while that of free Ubp6 is minimally affected (Fig. 2a, Extended Data Fig. 4a,b). The *ubp6-I329A L330A* mutant (hereafter *ubp6-AA*) exhibited nearly ideal behavior (Fig. 2b), with a stringent reduction of proteasome-activated deubiquitinating activity (to ∼2% of WT), and preservation of free activity. I329 and L330 are in close contact with Rpt1 in the structural model (Fig. 2c). Loss of deubiquitinating activity in the mutant could not be corrected by increasing the concentration of Ubp6; only a slight reduction of the affinity of Ubp6 for the proteasome was apparent (Extended Data Fig. 4c). Since Ub-AMC is not a true ubiquitin-protein conjugate, we tested Ubp6-AA on a ubiquitinated fragment of cyclin B^8^ (Ub_n_-NCB1). No deubiquitination was detected in the presence of Ubp6-AA (Fig. 2d). We also validated the *ubp6-AA* mutant by showing that it phenocopies *ubp6Δ in vivo* (Extended Data Fig. 4d,e).

**Fig. 4.**
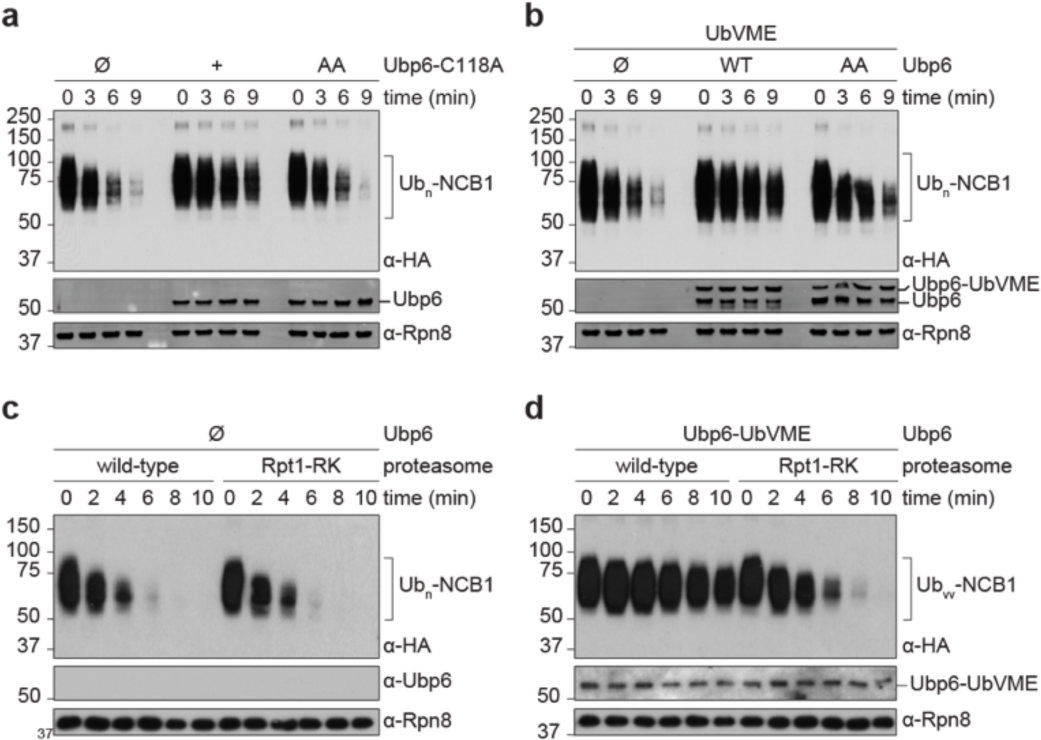
Ubp6-AA and Rpt1-RK mutants are impaired in the Ubp6 noncatalytic effect. **a**, *In vitro* degradation assay performed with C118A or the C118A I329A L330A triple mutant of Ubp6, proteasome, and HA-Ub_n_-NCB1 (detected with anti-HA antibody). Ubp6 and Rpn8 are loading controls. **b**, Inhibition of NCB1 degradation by Ubp6-UbVME. UbVME was pre-incubated with Ubp6 variants to covalently modify C118, abolishing deubiquitinating activity. The assay was otherwise as in *a*. **c**, NCB1 degradation by wild-type and mutant proteasomes. **d**, As *c* but with Ubp6-UbVME added.

To test whether the activation mechanism of Ubp6 is evolutionarily conserved, we generated substitutions in USP14 (human form), targeting the cognate residues of I329 and L330. Proteasome-dependent activation was reduced in the USP14-V343A, L344A mutant to ∼0.4% of wild-type, while the basal activity of free USP14 was unaffected (Fig. 2e and Extended Data Fig. 4f,g). Thus, the mutations identify a conserved mediator of Ubp6 activation, which we term the ILR element (Fig. 2f).

### Elements of the BL1 loop

Although the ILR element is within the BL1 loop, it is 10 residues removed from the segment of BL1 that occludes ubiquitin access to the active-site groove (Fig. 2f). How does the ILR reposition the inhibitory site of BL1? Figure 2c shows that the BL1 loop forms a previously unrecognized ß-hairpin (see Extended Data Fig. 3c for structural details). Ubiquitin is occluded by the ß8a strand of the hairpin; therefore activation should involve directed movement of this strand. The connection of ß8a to I329 and L330 is provided by ß8b, which is directly abutted by I329 and L330. Thus, we propose that the BL1 loop contains three distinct elements–the ILR, ß8a, and ß8b– which function in cooperation to control of Ubp6 activity (Fig. 2f). The sister strands of the BL1 loop are densely interconnected, so that the loop functions as a relatively rigid lever arm (Extended Data Fig. 3c,d) that efficiently propagates the allosteric signal emanating from the ILR element.

Repositioning of ß8a is expected to be insufficient to allow ubiquitin docking, as ubiquitin occlusion by the BL2 and SL would also have to be relieved (Extended Data Fig. 3a). However, unlike BL1, BL2 and SL do not contact the proteasome in our model (Fig. 1b). Examination of the crystal structure of free Ubp6 revealed that BL1 and BL2 are in direct contact, and SL is in contact with BL2, suggesting that transition of BL1 to the open form promotes the same change of state for the other blocking loops, as discussed below.

### Rpt1 mutant defective in Ubp6 activation

The Rpt1 components of Interfaces A and B lie within its OB domain (Fig. 1b), one of six proteasomal OB domains, which form a ring complex defining the substrate entry port of the RP^14^, but is to date not known to have any catalytic or regulatory function. To identify the proteasomal receptor site of the Ubp6 catalytic domain, OB domain mutants covering Interfaces A and B were generated. Several mutations impaired proteasome assembly and were not further studied (Fig. 3a and Extended Data Fig. 5a-c). Among assembly-proficient mutants, two were almost completely defective in activation of wild-type Ubp6: *rpt1-S164R T166K* and *rpt1-S164A T166K* (Fig. 3a). S164 and T166 are proximal to I329 and L330 of Ubp6 in our structural model (Fig. 3b). The *S164R T166K* double mutant (hereafter *rpt1-RK*), which has lost ∼97% of its capacity to activate Ubp6, was chosen for further analysis. S164 and T166 fall within a segment of the conserved L34 loop of the Rpt1 OB domain^14^ (Figs. 3c and Extended Data Fig. 5d), hereafter termed the activation loop.

**Fig. 5.**
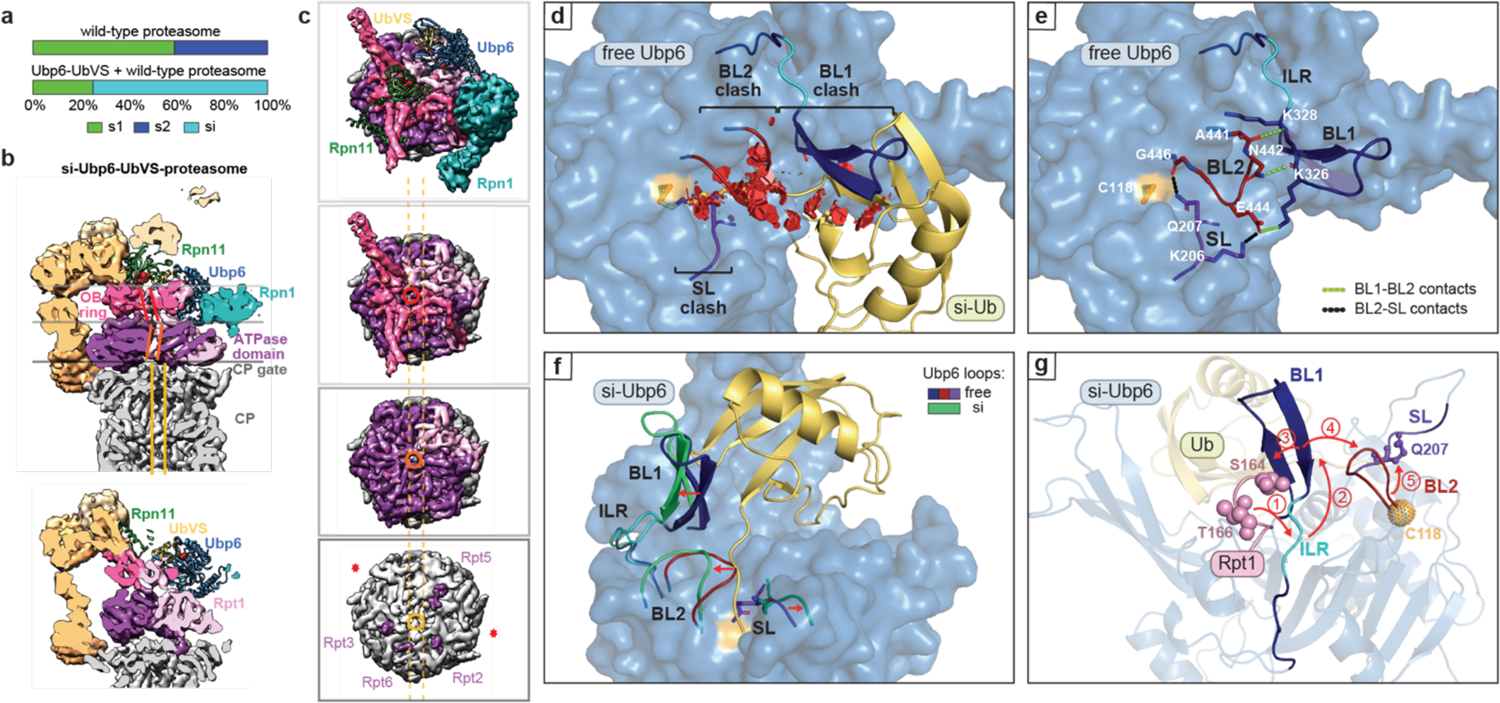
The inhibitory Ubp6-UbVS-proteasome complex defines a new conformational state of the proteasome. **a**, Ubp6-UbVS induces the si proteasome state. **b**, Cross-section of the cryo-EM density map of the si-state proteasome. Misaligned axial channels of the OB ring, the ATPase domain ring, and the CP are denoted by parallel bars (red, orange, and yellow, respectively). Rpn11 (green) is misaligned with the OB domain substrate entry port. Horizontal lines indicate cutting planes in ***c***. Bottom: alternative z-plane visualizes Ubp6 catalytic domain. Active sites of Ubp6 and Rpn11 are highlighted in red, proteasome subunits as in Fig. 1a. **c**, Cut-away views down the long axis of si proteasome from the OB ring to the ATPase domain ring to the CP α-ring. Subunits of interest are highlighted. RP axial channels are off-axis to the CP (dotted yellow lines). The OB substrate entry port is circled in red, ATPase translocation channel in orange. Four C-terminal Rpt tails (Rpt2, Rpt6, Rpt3, Rpt5) are inserted into CP α pockets. Asterisks: unoccupied α pockets. **d**, Occlusion of ubiquitin by BL1, BL2, and SL in free Ubp6 illustrated by red discs (PDB: 1VJV). Ubiquitin is modelled onto free Ubp6, positioned as in complex si. **e**, BL1-BL2-SL network in free Ubp6 (PDB: 1VJV) is established through interloop contacts. **f**, Comparison of the blocking loop network in free Ubp6 (PDB: 1VJV) and complex si. Loops from free Ubp6 are superimposed onto si-Ubp6. Structure is rotated 90° counterclockwise from *d*. **g**, Proposed cascade of signal transfer within Ubp6 upon interaction with proteasome.

Rpt1-RK proteasomes appeared to be inherently unable to activate Ubp6, and not simply attenuated in Ubp6-proteasome affinity, as the catalytic defect could not be overcome by adding elevated levels of Ubp6 to the reaction (Fig. 3d, Extended Data Fig. 5e). The failure in deubiquitination extended to bona fide ubiquitin-protein conjugates (Fig. 3e) and was confirmed by *in vivo* assays (Extended Data Fig. 5f,g).

### Relief of proteasome inhibition

When the catalytic cysteine of Ubp6 is substituted with alanine (Ubp6-C118A), the resulting enzymatically inactive protein still inhibits the proteasome^15^. This “noncatalytic effect” is not an aberrant feature of the mutant, but an inherent property of Ubp6, since it is also seen with wild-type Ubp6 when it is coupled to ubiquitin in adducts such as Ubp6-UbVS^15, 16^. Are mutants in which the proteasome cannot activate Ubp6 also defective in proteasome inhibition? To test this possibility, *in vitro* degradation assays were performed using Ub_n_-NCB1 as substrate. We observed a strong noncatalytic effect when degradation assays were performed in the presence of Ubp6-C118A (Fig. 4a). The effect was almost completely abrogated by the Ubp6-C118A-AA triple mutant protein. Failure of the noncatalytic effect could simply result from deficient ubiquitin engagement by this mutant. However, when we used Ubp6-AA, modified covalently at Cys118 by UbVME, we similarly observed a strong impairment of the noncatalytic effect (Fig. 4b). Thus, even when ubiquitin occupies the active site of Ubp6, forcing the blocking loops open, the Ubp6-AA mutant protein cannot inhibit the proteasome.

The Rpt1-RK mutant proteasome was indistinguishable from wild-type when tested in a degradation assay with Ub_n_-NCB1 as substrate and no Ubp6 present (Fig. 4c), exemplifying the specific nature of this mutant. However, the mutant restored substrate degradation in the presence of Ubp6 (Fig. 4d). Thus, the L34 activation loop serves as the receptor element in the proteasome for the noncatalytic effect exerted by Ubp6. In both activation of Ubp6-mediated deubiquitination and the noncatalytic effect, the *rpt1-RK* mutant phenocopies *ubp6-AA*. Abrogation of the noncatalytic effect was also shown *in vivo* for both mutants (Extended Data Fig. 6). In summary, these results indicate that Ubp6 activation and proteasomal inhibition are inherently coupled processes.

### Structure of Ubp6-inhibited proteasomes

In the proteasome-Ubp6-UbVS ternary complex, the catalytic domain of Ubp6 docked at Rpt1 and exerted a dramatic influence on the structure of the proteasome, driving 75% of proteasomes into a novel conformational state that we term si (Fig. 5a). Proteasomes in the basal state, s1, exhibit axial misalignment–a signature of their inactivity. This is also true of proteasomes in the s2 and s5 states^17–19^. With substrate engagement and conversion to an active state such as s3 or s4, co-axial positions are assumed by the active site of Rpn11, the substrate entry port of the OB ring, the central channel of the ring formed by the six ATPase domains, and the heptameric α ring of the CP. These structural elements are misaligned in si proteasomes (Fig 5b,c). Also indicative of an inactive state is the closed gate of the CP (Fig. 5c). The positioning of the lid of si resembles that of s5^17^, whereas the ATPase ring is comparable to that of s2, except that the C-terminal tail of the Rpt6 is inserted into the α2/α3 pocket of the CP α ring, as seen in s3 (Extended Data Fig. 7). In summary, si proteasomes borrow features from a variety of other states to form a unique and degradation-inhibited conformational state.

In free Ubp6, BL1 is stabilized at its base on opposite sides by the PKL and BL3 loops, which notably both contact the ILR element (Extended Data Fig. 3e). In contrast, the PKL and BL3 loops move away from the ILR element in the si complex, which may facilitate contact with the activation loop of Rpt1 and movement of the ILR element (Extended Data Fig. 3f).

Cryo-EM analysis of the Rpt1-RK proteasome together with wild-type Ubp6 and UbVS revealed that the fraction of proteasomes in the si state was reduced to ∼25%, whereas the conformational profile of the Rpt1-RK proteasome alone was comparable to that of wild-type proteasomes (Extended Data Fig. 8a). Thus, the Rpt1 mutation impairs the Ubp6-dependent transition of the proteasome to the si state, although Ubp6-UbVS remains docked at Rpt1 in the s2 state proteasomes that are observed with the mutant (Extended Data Figs. 1e,7d). Release of the proteasome from the si state may account for the recovery of protein degradation by the mutant (Fig. 4). The mutation also conferred structural changes on the associated Ubp6 enzyme: Ubp6 was slightly shifted from its position on wild-type proteasomes, and the activation loop retracted from the ILR (Extended Data Fig. 8b-d). In summary, a modest perturbation of Rpt1-ILR contact interface can decisively alter the conformational profile of the proteasome as a whole.

### The suppressive network of Ubp6

Comparison of the structure of Ubp6 associated with si proteasomes to the 1.7Å structure of free Ubp6 provided major insights into the mechanism of Ubp6 activation. BL1, BL2, and SL are all in position to clash with ubiquitin in free Ubp6 and must be displaced to activate the enzyme (Fig. 5d). In free Ubp6, BL1 directly contacts BL2 through three hydrogen bonds, extending from ß8b, directly adjacent to the ILR element (Fig. 5e). BL2 in turn contacts SL through a salt bridge; while G446, immediately flanking BL2, directly contacts the side chain of Q207, which is the key ubiquitin-blocking residue of SL (Fig. 5d,e and Extended Data Fig. 3a). Thus, BL1, BL2, and SL stabilize each other to form an inhibitory network of blocking elements, accounting for the tight suppression of activity in free Ubp6. In the activated Ubp6 of the si structure, all three blocking loops are withdrawn in from the catalytic groove (Fig. 5f, Extended Data Fig. 9). Proteasome contact with the ILR may direct repositioning of the ß-hairpin, with movements propagated in an ordered sequence across the network of blocking loops (Fig. 5g). The direct target of ILR movement is ß8b, opposite sides of which are in contact with ubiquitin-blocking elements ß8a and BL2.

## Discussion

We have identified an allosteric network composed of the activation loop of Rpt1; its target, the ILR element of Ubp6; and downstream elements, the BL1 ß-hairpin, the BL2 loop, and the SL loop. We propose that the signal generated by Rpt1-Ubp6 interaction is propagated stepwise across these elements, in a progression initiating at Rpt1 and ending at the SL loop. Network components exert control over both Ubp6 and the proteasome, activating the former and inhibiting the latter (Extended Data Fig. 10). This coupling is associated with the si state of the proteasome, which is induced by the ILR allosteric switch. The pausing of proteasome-mediated substrate degradation in the si state may impose temporal order on otherwise competing enzymatic reactions, and provide an extended, substrate-controlled time window for Ubp6 to remove ubiquitin groups.

Our findings have general implications for the regulation of deubiquitinating enzymes of the 56-member USP family^20^. Since promiscuous deubiquitination by these enzymes has the potential to neutralize the myriad functions of ubiquitination, it is essential that their activities are held under negative control and allosterically activated with specificity at a given time or location. Crystallographic studies from the Shi lab^13, 21^ identified blocking loops in these enzymes, BL1 and BL2, and hypothesized that they may be involved in control of activity. This idea remained hypothetical, and, surprisingly, subsequent studies have instead favored the view that blocking of the catalytic cleft by BL1 and BL2 is not critical for suppression per se. In this interpretation, BL1 and BL2 are found in an open state in substrate-engaged forms of USP enzymes, such as UbVS-modified USPs, simply by virtue of substrate accommodation^22–25^. However, the blocking loops are primordial features of these enzymes, and the concept that they move primarily or only by substrate accommodation has not been adequately reconciled with their conservation over the eukaryotic kingdom and the USP family as a whole. Thus, our finding that these loops are central to allosteric control in Ubp6 suggests an important paradigm for this enzyme family.

We have found that BL1, BL2, and SL operate as an integrated network, with BL1 serving as the fulcrum for rearrangement of BL2 and SL. The participation of three distinct loops in blocking the catalytic cleft may ensure tight negative control. In USP enzymes where the evolutionary pressure to repress basal activity is less strong, some of the loops may have degenerated to a more flexible and less repressive state. Thus, while Ubp6 may provide a clear-cut case of blocking loop action because of the strength of the allosteric control mechanism, owing to the highly defined closed state of its blocking loops, the essential features of the mechanism are likely to apply to many enzymes of the USP family.

## Methods

### Expression and purification of recombinant proteins

Unless otherwise noted, Rosetta (DE3) cells (EMD Millipore) transformed with expression plasmids were grown to an OD_600_ of 0.6-0.8 in selective 2X YTG media (10 g/L yeast extract, 16 g/L tryptone, 20 g/L dextrose). Expression was induced with 0.5 mM IPTG (Gold Biotech), and cells were transferred to a 16°C shaker for overnight induction. Cells were harvested by centrifugation, and purification was carried out as described below. All lysis buffers used were supplemented with 1X protease inhibitor cocktail^11^ and 2 mM AEBSF (Gold Biotech). A list of constructs used can be found in Supplementary Table 2.

### His6-tagged Ubp6 proteins

Cell pellets were resuspended with His-tag lysis buffer (50 mM NaH_2_PO_4_, 100 mM NaCl, 10% glycerol [v/v], 25 mM imidazole [pH 8.0]). Cells were lysed by French press at 10,000 psi (two passes), and the lysate was clarified by centrifugation at 20,000 x *g* for 30 min at 4°C. 2 mL Ni-NTA resin (Qiagen) was used for clarified cell lysate from 1 L bacterial culture. Resin was incubated with clarified lysate for 2 h at 4°C, followed by washing with 80 bed vol of wash buffer (50 mM NaH_2_PO_4_, 300 mM NaCl, 10% glycerol [v/v], 25 mM imidazole, [pH 8.0]). Stepwise elution was achieved using a total of 10 mL elution buffer (50 mM NaH_2_PO_4_, 100 mM NaCl, 10% glycerol [v/v], 250 mM imidazole [pH 8.0]). Peak fractions containing significant amounts of protein were pooled. All Ubp6 variants were prepared using His6-tagged constructs.

### Generation of Ubp6-UbVS and Ubp6-UbVME adduct

A 5-fold molar excess of ubiquitin-vinyl-sulfone (UbVS) or ubiquitin-vinyl-methyl-ester (UbVME) was incubated with different Ubp6 variants in reaction buffer (50 mM Tris-HCl [pH 7.5], 1 mM DTT) for 2.5 h at 30°C. Adducts were purified by FPLC on a Superdex 200 HiLoad 16/600 column (GE Healthcare) to remove unmodified Ubp6. Purified adducts were stored in a buffer of 20 mM Tris-HCl (pH 7.5), 50 mM NaCl at −80°C.

### GST-tagged USP14

Pelleted cells were resuspended and lysed in PBS (137 mM NaCl, 2.7 mM KCl, 10 mM Na_2_HPO_4_, 1.8 mM KH_2_PO_4_ [pH 7.4]). 500 μL Glutathione Sepharose 4B resin (GE Healthcare) was used for clarified cell lysate from 1 L bacterial culture. Resin was incubated with clarified lysate for 2 h at 4°C, followed by washing with 50 bed vol of PBS, then 50 bed vol of PBS with 100 mM NaCl, and lastly with 50 bed vol of PBS. To remove the GST tag, resin was incubated with 2 bed vol of PBS containing 10 μL of 1 U/μL thrombin (Sigma) for 2 h at 25°C with occasional agitation. To remove thrombin, 100 μL Benzamidine-Sepharose (GE Healthcare) was then added, and the eluate was incubated for 30 min at 4°C with rocking. Glycerol was added to the eluates at 10% (v/v) final concentration for storage at −80°C.

### Recombinant proteasome base for HXMS experiments

The three plasmids used for recombinant expression of yeast base subcomplex were a kind gift from Dr. A. Martin (UC Berkeley): pCOLADuet-Rpt1-Flag, His6-Rpt3, Rpt2, Rpt4-6 (kanamycin), pETDuet-Rpn1,2,13 (ampicillin) and pACYCDuet-Nas2, Nas6, Hsm3, Rpn14 (chloramphenicol), and tandem affinity purification was carried out as described^26^.

### Purification of yeast or human proteasomes

#### Purification of 26S yeast holoenzyme and proteasome subcomplexes

Protein A-tagged 26S proteasome and regulatory particle (RP) used for biochemical assays were affinity-purified as described^27^, and yeast strains used are listed in Supplementary Table 3. Purification of wild-type and Rpt1-RK mutant 26S proteasomes with a 3X FLAG tag used for cryo-electron microscopy studies was performed as described^18^, and yeast strains used for this purpose are listed in Supplementary Table 4.

### Purification of biotin-tagged human proteasome

Human proteasome holoenzyme was purified via affinity tag as previously described^8^. To eliminate UCH-L5 activity, human proteasomes were treated with ubiquitin-vinyl-sulfone (UbVS, Boston Biochem). UbVS was added to the resin at 1-1.5 mM, followed by incubation at 30°C for 2 h prior to TEV cleavage. Residual UbVS was removed by washing the resin with at least 20 bed vol of low-salt buffer. The Ub-AMC hydrolysis assay was used to confirm the elimination of UCH-L5 activity. The purity and integrity of the purified proteasomes were routinely assessed using native gels^28^ and SDS-PAGE.

### Cryo-electron microscopy (cryo-EM) studies

#### Sample preparation

To study the structure of the 26S-Ubp6-UbVS complex, 26S proteasomes and Ubp6-UbVS were mixed in a 1:4 ratio and incubated for 20 min on ice before plunging in a solution containing 20 mM HEPES-NaOH (pH 7.4), 40 mM NaCl, 4 mM DTT, 4 mM MgCl_2_, 4 mM ATP, and ∼25% sucrose. Cryo-EM data of the plunged samples, 26S-Ubp6-UbVS (400 nM 26S, 1.6 μM Ubp6-UbVS), free 26S Rpt1-RK (400 nM), 26S Rpt1-RK-Ubp6-UbVS (400 nM 26S Rpt1-RK, 1.6 μM Ubp6-UbVS) were collected on a Titan Krios (Thermo Fisher) with a K2 or K3 detector (Gatan Inc.). Images were acquired in counting mode at a pixel size of 1.09 Å for K3 camera and 1.38 Å for K2 camera (Supplementary Table 1). Each total exposure of 60 electrons per Å^2^ was fractionated into 30 frames for K3 camera, while 35 electrons per Å^2^ into 33 frames for K2 camera. Defocus ranged from −1.0 to −2.5 µm (K3) and −1.8 to 3.0 µm (K2).

### Data processing

Initial motion correction was done by MotionCor^29^ as implemented by RELION 3.0^30^. Contrast transfer function estimation was performed by CTFFIND4^31^. Particles were picked either by REION or Cryolo^32^. The following processing was done in RELION 3.0, unless otherwise specified. Two rounds of 2D classification and one round of 3D classification were performed to enable selection of double-capped particles for further processing. A published map (EMD-3534) was low-pass filtered to 60 Å and used as a reference for initial 3D classification. A C2 symmetry expansion was performed, followed by subtraction of a single 19S cap density from the double-capped particles. A refinement applying C2 symmetry and Bayesian polishing improved the resolution of the maps. Final classes were compared with known conformations and assigned to the conformational states. To improve the resolution around Ubp6 in the si^WT^ structure, a 2-body refinement was performed with the si^WT^ structure, separating the flexible Rpn1 and ATPase density (body2) from the rest of the single-capped proteasome (body1). All maps were sharpened by phenix.auto sharpen^33^.

### Model construction of Ubp6 bound proteasome states in cryo-EM analysis

#### Initial Ubp6 model with bound ubiquitin-vinyl-sulfone

We first constructed a complete model of the catalytic (CAT) domain of Ubp6 (residues 104 to 499) based on the 1.74Å resolution yeast Ubp6 crystal structure with the PDB: 1VJV. Structural elements not resolved in the crystal structure (residues 200 to 203, 174 to 177, 283 to 293, and 370 to 387) were modeled using the Rosetta^34^ *ab initio* structure prediction framework implemented as plugin in VMD 1.9.4a35 software^35^. This framework was initially developed to furnish the structurally unresolved regions of the 26S proteasome^36^. For each insertion larger than three residues, the predicted 5000 models for each of the domains where clustered using the partitioning around medoids cluster algorithm. The representative structure with the best score was used to complete the unresolved domains. The resulting completed model of the CAT domain of Ubp6 was then docked as rigid body into the cryo-EM density using Chimera^37^.

In all states, we identified an extra density where ubiquitin usually binds to Ubp6. So we constructed an Ubp6-CAT complex with bound ubiquitin. The human ubiquitin-aldehyde bound USP14 structure (PDB: 2AYO) was aligned with the obtained Ubp6-CAT model. Then the coordinates of the ubiquitin-aldehyde chain (chain ID B) of the aligned 2AYO crystal structure were pasted into our Ubp6-CAT model. The resulting Ubp6-CAT-ubiquitin-vinyl-sulfone (Ubp6-CAT-UbVS) model was then docked as rigid body into the remaining cryo-EM densities.

### Initial 26S proteasome models

To model the CP and the lid of the 26S yeast proteasome, we used the structures of the different proteasome states without bound Ubp6 from Eisele et al.^37^ as initial structures for rigid body docking. The exact subunits used for each state are detailed in Supplementary Table 1. The Rpt1 mutants were generated using the mutagenesis plugin in VMD to replace in Rpt1 serine 164 by an arginine and threonine 166 by a lysine.

### Model fitting into the cryo-EM maps

The aforementioned structures for each of the states were fitted into the respective cryo-EM map using molecular dynamics flexible fitting (MDFF)^17^. MDFF employs molecular dynamics to fit initial models into a density in real space, and thus permits protein flexibility while maintaining realistic protein conformations^38^. We used NAMD^39^ with the CHARMM36 force field for MDFF calculations. During MDFF runs, restraints to preserve the secondary structure, chirality, and cis-peptide bonds were applied to avoid overfitting. As further step to reduce artifacts due to overfitting, all MDFF runs were performed at a modest gscale of 0.3.

First the Ubp6-CAT models and if available the Ubp6-CAT-UbVS interface were fitted into the density with MDFF while fixing the rest of the 26S proteasome. We employed an additional constraint between the catalytic Cys118 of Ubp6-CAT and the C-terminus of UbVS. Such constraint is necessary, as the catalytic cysteine is trapped by UbVS in a thioester bond formed between the C-terminus of ubiquitin and Cys118, but parametrization of this type of bond is not available in the CHARMM36 force field.

In a subsequent MDFF run, the whole structure (including all 26S proteasome subunits) was refined. Each of these runs started with 200 steps of energy minimization followed by 40ps MDFF simulation at a temperature of 300K. The Ubp6-CAT-Rpt1 and Ubp6-CAT-UbVS interfaces were further refined using interactive MDFF to manually pull side chains to the desired regions of density, while interactively checking the cross-correlation values in VMD. Interactive MDFF runs were performed using the MDFF graphical user interface and initiated using QwikMD routines^40^.

Finally, we checked the hydrogen bond network with the reduce^41^ module of Phenix software^42^ and performed Phenix real space refinement with reference coordinate restraints (σ=0.05) on the whole structure. We removed hydrogen atoms for model deposition in the PDB as hydrogen atoms are not resolved in any of the density maps.

### Structural data analysis

Structural models and density maps were analyzed and visualized using VMD, Chimera, PyMol, and Coot^43^. Interaction patterns were identified using PyContact^44^ and the contact matrix algorithm implemented in Maximoby (CHEOPS, Germany). The structures were validated using MolProbity^45^. The results of the structure validation are detailed in Supplementary Table 1.

### Hydrogen-deuterium exchange mass spectrometry (HXMS)

#### Deuterium labeling

To monitor exchange in the RP, it was incubated alone or with a 5-fold molar excess of Ubp6-UbVME for 1 h on ice to allow for complex formation. To monitor exchange in Ubp6 and Ubp6-UbVME, His6-Ubp6 or His6-Ubp6-UbVME were incubated alone or with a 1.65-fold molar excess of proteasome base for 1 h on ice to allow for complex formation. After incubation, the complexes were diluted 12-fold with labeling buffer (10 mM HEPES-NaOD [pD 7.5] 50 mM NaCl, 50 mM KCl, 5 mM MgCl_2_, 0.5 mM EDTA, 0.5 mM ATP, 1 mM DTT, 10% glycerol [v/v], D_2_O [Cambridge Isotope Laboratories]) at 25°C and quenched by a 2-fold dilution with ice-cold quench buffer (0.8 M guanidine hydrochloride, 0.8% formic acid [v/v], H_2_O) at time points ranging from 10 s to 4 h. Undeuterated control samples were prepared for each of the proteins alone and in complexes using the same procedure as outlined above and with buffer made using H_2_O instead of D_2_O.

### Liquid Chromatography and Mass Spectrometry (LC/MS)

Samples were immediately digested offline after the addition of quench buffer with 10 μL of a 50% (v/v) slurry of immobilized porcine pepsin (prepared in house using POROS beads) for 5 min on ice. The remaining workflow and analysis have been previously described^11^, except that peptides were eluted and separated using a 5%–35% gradient of acetonitrile over 18 min. The M-class Acquity UPLC with HDX technology coupled to a Waters Synapt G2-Si HDMS^E^ mass spectrometer was used for analysis and calibrated with direct infusion of a solution of glu-fibrinopeptide (Sigma) at 200 fmol/uL prior to data collection. A conventional electrospray source was used with a temperature of 80°C and desolvation temperature of 175°C. All comparison experiments were done under identical experimental conditions such that deuterium levels were not corrected for back-exchange and are therefore reported as relative.

### Data processing

Peptides were identified using PLGS 3.0.1 (Waters, RRID: SCR_016664, 720001408EN) using multiple replicates of undeuterated control samples. Raw MS data were imported into DynamX 3.0 (Waters, 720005145EN) and filtered as shown in Supplementary Table 5. Those peptides meeting the filtering criteria were further processed automatically by DynamX followed by manual inspection of all processing. The relative amount of deuterium in each peptide was determined by subtracting the centroid mass of the undeuterated form of each peptide from the deuterated form, at each time point, for each condition.

### Ub-AMC hydrolysis assays

All reactions were performed in Ub-AMC assay buffer (50 mM Tris-HCl [pH 7.5], 1 mM EDTA, 1 mM ATP, 5 mM MgCl_2_, 1 mM DTT, and 1 mg/mL ovalbumin [Sigma]), with a final reaction vol of 20 μL per assay in a 384-well plate (Corning). Ub-AMC cleavage was monitored by measuring fluorescence in real time for at least 30 min at 365 nm excitation and 460 nm emission with an EnVision plate reader (Perkin Elmer, ICCB Facility, HMS). The initial kinetics observed within the linear range was used for plotting. For routine assays, Ub-AMC (Boston Biochem) was used at a concentration (0.5 to 1 μM) far below the K_M_, so that activity measurements were performed under k_cat_/K_M_ conditions. All Ub-AMC assays were independently repeated and yielded consistent results.

### Ubp6 activity in mutagenesis screens

For free Ubp6 activity, Ubp6 variants were assayed at 0.5 μM in Ub-AMC assay buffer. The reaction was initiated by adding Ub-AMC to 0.5 uM. For proteasome-bound Ubp6 activity, Ubp6 variants (4 nM final) were pre-incubated with *ubp6Δ hul5Δ* yeast proteasome (1 nM final) in Ub-AMC assay buffer at 25°C for 15 min. The reaction was initiated by adding Ub-AMC to 0.5 μM.

### Measurement of Ubp6 affinity to proteasome

To estimate the affinity of Ubp6-AA for wild-type proteasome, and the affinity of wild-type Ubp6 for Rpt1-RK proteasome; the activation of Ubp6 for Ub-AMC hydrolysis was used as a proxy for association. Assays were carried out using Ubp6 concentrations ranging from 1.25 nM to 60 nM. 1 μM Ub-AMC and 1 nM wild-type and *rpt1*-mutated *ubp6Δ hul5Δ* yeast proteasome were used. Hydrolysis rates of Ubp6 alone and proteasome alone were subtracted from the rates observed for Ubp6 in the presence of the proteasome. The Ubp6 concentration and rate data were fit to a hyperbolic curve by nonlinear regression using the PRISM software.

### USP14 activity assay

For free activity, USP14 variants were assayed at 1 μM in Ub-AMC assay buffer. The reaction was initiated by adding Ub-AMC to 1.5 μM. For proteasome-bound activity, USP14 variants (8 nM final) were pre-incubated with UbVS-treated human proteasome (1 nM final) in Ub-AMC assay buffer at 25°C for 15 min. The reaction was initiated by adding Ub-AMC to 1 μM.

### Preparation of APC/C-mediated ubiquitinated conjugates

APC/C was immunopurified from *X. laevis* egg extract with anti-Cdc27 antibody (Santa Cruz) bound to Protein-A Sepharose (Sigma), and ubiquitination reactions were performed as described^8, 46^. Alternatively, Strep-tagged APC and its activator His-Cdh1 were purified from insect cells infected with baculovirus-based expression vectors^47^. In the latter case, 0.015 μM Strep-APC and 0.3 μM His-Cdh1 were first preincubated on ice in 1X TBS (25 mM Tris-HCl [pH 7.5], 50 mM NaCl) for 30 min to activate the APC. The conjugation mixture contained 0.3 μM MBP-E1, 1.5 μM His-UbcH10, 0.13 μM His-Ube2S, 100 μM ubiquitin, 5 μM HA-tagged N-terminus fragment of cyclin B1 (HA-NCB1), and the activated APC/C. A 100 μL reaction was typically performed in 1X reaction buffer (25 mM Tris-HCl [pH 7.5], 50 mM NaCl, 10 mM MgCl_2_) in the presence of ATP regeneration system (3.5 U/mL creatine kinase [Roche], 3.8 mM creatine phosphate disodium [Roche], 0.5 mM ATP disodium salt, 0.5 mM MgCl_2_) for 5 h at 25°C. Glycerol was added to the ubiquitinated NCB1 conjugates (HA-Ub_n_-NCB1) to a final concentration of 10% (v/v) for storage at −80°C.

### *In vitro* degradation and deubiquitination assays

Purified recombinant Ubp6 variants (Fig. 4a-b: 120 nM; Fig. 4c-d: 240 nM) were first incubated with nominally wild-type (Fig. 4a-d: *ubp6Δ hul5Δ* ecm29*Δ*, 30 nM) or mutant (Fig. 4c-d: *rpt1-RK ubp6Δ hul5Δ* ecm29*Δ*, 30 nM) yeast proteasomes for 15 min. Where indicated, Ubp6 was pre-incubated with 5-10-fold excess of UbVME to induce adduct formation prior to incubation with the proteasome. 200 nM HA-Ub_n_-NCB1 was then added to the Ubp6-proteasome mixture to initiate the reaction. At the indicated time points, 10 μL of the reaction mixture was withdrawn, and the reaction was terminated by adding 2X Laemmli loading buffer and heating at 70°C for 10 min. Samples were subsequently analyzed by SDS-PAGE and immunoblotting.

For deubiquitination assays uncoupled to degradation, ADP, proteasome and HA-Ub_n_-NCB1 were first rendered ‘ATP-free’ by treating with hexokinase and glucose as previously described^8^. Purified recombinant Ubp6 variants (120 nM) and ADP-proteasomes (Fig. 2d, 3e: *ubp6Δ hul5Δ* i.e. nominally wild-type; Fig. 3e: *rpt1-RK ubp6Δ hul5Δ*, 5 nM) were assayed in reaction buffer (50 mM Tris-HCl [pH 7.5], 5 mM MgCl_2_, 0.5 mM DTT) supplemented with 5 mM ‘ATP-free’ ADP. Proteasome and metalloprotease inhibitors were also supplemented in the reaction, which includes 0.5 mM ATPγS (Santa Cruz), 1.5 μM PS-341 (ApexBio), 7.5 μM MG-262 (Apexbio), 100 μM Epoxomicin (ApexBio), and 10 mM 1,10-phenanthroline (o-PA, Sigma). Ubp6 and ADP-proteasomes were preincubated for 15 min at 30°C. 200 nM ADP-HA-Ub_n_-NCB1 was then added to the Ubp6-proteasome mixture to initiate the reaction. Samples were collected and analyzed as above. All *in vitro* assays have been independently repeated. Yeast strains used are listed in Supplementary Table 3, and antibody information can be found in Supplementary Table 6.

### Yeast methods and media

Standard techniques were used for strain constructions and transformations. A list of plasmids used for yeast transformation can be found in Supplementary Table 7. For plate assays, strains were inoculated into either YPD (1% bacto-yeast extract, 2% bacto-peptone, 2% dextrose, 50 mg/L adenine, 400 mg/L tryptophan, 500 mg/L uridine) or selective media and grown overnight at 30°C. Cell density was measured, and the cultures were diluted with fresh media to OD_600_= 0.1. Cultures were allowed to grow at 30°C for another 3-5 h, until the OD_600_ reached log phase. Cultures were then serially diluted three-fold in media, spotted onto plates with a pin array, and incubated at 30°C for 3-7 days.

All plates were prepared using media supplemented with 2% agar. Synthetic plating medium consisted of 2% dextrose, 0.67% yeast nitrogen base without amino acids, 12.5 mg/L adenine hemisulfate, 125 mg/L uridine, 50 mg/L phenylalanine, 50 mg/L isoleucine, 75 mg/L valine, 10 mg/L tyrosine, 150 mg/L proline, 25 mL/L glycine, 75 mg/L alanine, 75 mg/L serine, 50 mg/L threonine, 100 mg/L glutamate, 50 mg/L aspartate, 200 mg/L glutamine, 50 mg/L asparagine, 37.5 mg/L histidine, 112.5 mg/L lysine, 37.5 mg/L methionine, 100 mg/L arginine, 100 mg/L leucine, 100 mg/L tryptophan. Certain amino acids were excluded from the mixture according to the test plate formulation.

For the Ub-K-Trp experiment, medium without uridine (2% dextrose, 0.67% yeast nitrogen base without amino acids, 0.5% casamino acids, 50 mg/L adenine hemisulfate, 500 mg/L tryptophan) was used in Extended Data Fig. 4d to select for the transforming plasmid carrying Ubp6 variants with *URA3* marker. Medium in Extended Data Fig. 5e was prepared similarly, except with the addition of 500 g/L uridine, as these are uracil auxotrophs. Yeast strains used for Ub-K-Trp assay can be found in Supplementary Table 8.

For canavanine experiments (Extended Data Figs. 4e and 5g), arginine was omitted because it competes with canavanine. For Extended Data Fig. 4e, uridine and tryptophan were additionally omitted (Ubp6 variants with *URA3* maker and ubiquitin overexpression or corresponding control plasmid with *TRP1* marker) where needed to maintain selection. Canavanine was added to a final concentration of 5 μg/mL. For plasmids in which ubiquitin expression is driven by the *CUP1* promoter, media were supplemented with 100 μM copper sulfate. Yeast strains used for canavanine assays can be found in Supplementary Table 9.

At least five transformants were picked 3-4 days post-plating. Each set of plate assays was done with at least three transformants and independently repeated, all yielding equivalent results. Representative images were shown in this study.

For evaluating the Ubp6 noncatalytic effect *in vivo*, a colony forming assay was used (Extended Data Fig. 6). Plasmids expressing variants of Ubp6 were transformed into either *ubp6Δrpn4Δ* (sJH185) or *ubp6Δ* (sJH183) yeast strains. Colony formation was scored after incubation at 30°C for 3-4 days. *RPN4* deletion eliminates the cells’ capability to counteract proteasome stress by upregulating proteasome subunit synthesis. The presence of catalytically inactive *ubp6-C118A* mutation results in cell death due to ubiquitin stress (i.e., ubiquitin depletion resulting from failure of catalytic deubiquitination prior to substrate degradation), together with proteasome stress from degradation inhibition (i.e., the noncatalytic effect). This assay has been independently repeated and yeast strains used for this assay can be found in Supplementary Table 10.

### Reagents

Additional information on commercial reagents used in this study is detailed in Supplementary Table 11.

## Data and materials availability

Plasmids and yeast strains are available upon request. EM maps and models are available through EM Data Bank (EMDB) and the Protein Data Bank (PDB) under the following accession codes:

## Acknowledgements

We thank W. Baumeister and J. Plitzko for providing support and cryo-electron microscopy instrumentation; M. Hartwig (MPI Biochemistry) for sample preparation; Y. Tone, D. Schibich, A. Schwarz, and A. Ulmer for contributing to yeast strain constructions; and T. Sixma and E. Strieter for critical reading of the manuscript. This research was funded by grants from the National Institutes of Health (R01 GM043601 to D.F., R01-GM101135 to J.R.E, R01-GM134064-01 to Y.L.), by a Dean’s initiative grant to D.F. and Y.L., and by a research collaboration with the Waters Corporation (J.R.E.). This work was also supported by the Deutsche Forschungsgemeinschaft (DFG) through Germany’s Excellence Strategy - EXC 2067/1-390729940, SFB1035/Project A01, and CRC889/Project A11, and Marie Curie Career Integration grant (PCIG14-GA-2013-631577) to E.S., by the Gauss Centre for Supercomputing e.V. for providing computing time on the GCS Supercomputer SuperMUC at Leibniz Supercomputing Centre (project id: pn56ri) to S.K. and T.R., and by the National Research Foundation of Korea (NRF) grants and the DGIST R&D program of the Ministry of Science and ICT of Korea (2019R1A4A1024278, 2019R1A2C1089413, & 21-CoE-BT-04) to B.-H.L.

## Author contributions

Biochemical studies and *in vivo* experiments were performed and analyzed by K.Y.S.H., S.S, S.E., T.S. with guidance from S.E., G.T., B.-H.L., X.-H. F., and D.F.; cryo-EM studies and structural modeling were conducted by S.K., M.R.E., T.C.C., T.J., T.R., G.T., S.E., and E.S.; hydrogen deuterium mass spectrometry experiments were performed by J.A.M. under the supervision of J.E.; UbVME was generated by D.V.D. and H.O.; ubiquitin conjugates were prepared by K.Y.S.H., B.-H.L. and Y.L.; the paper was written by D.F., K.Y.S.H., T.R., and E.S.; and was critically reviewed by all authors.

## Additional information

### Supplementary information

**Correspondence and requests for materials** should be addressed to D.F., E.S., T.R., and B.-H.L.

#### Supplementary Discussion

Our model for Ubp6 activation on the proteasome incorporates previous findings and highlights the varied roles played by ubiquitin and ubiquitin-like (UBL) domains in the process (Extended Data Fig. 10). In the absence of its N-terminal UBL domain, Ubp6 exhibits only basal activity *in vitro*, and the UBL deletion behaves as a null mutation *in vivo*^10^. The UBL docks at the T2 site of Rpn1, mutation of which also phenocopies a *ubp6* null^11^. Thus, assembly of the catalytic complex is proposed to begin with the UBL docking at T2 (Extended Data Fig. 10, step 1). The interaction between Ubp6 and Rpn1 appears to promote Ubp6-Rpt1 interaction by increasing the local concentration of the Ubp6 catalytic domain as well as orienting the domain towards Rpt1.

The following steps entail displacement of the blocking loops of Ubp6 (Extended Data Fig. 10). In this model, the loops are first partially displaced in a hypothetical priming reaction that will be discussed more fully elsewhere. Partial blocking loop displacement enables target ubiquitin (T) to dock at the Ubp6 active site. In step 3, the proteolytic substrate is docked on the proteasome, though not to Ubp6; initial docking involves a second ubiquitin on the conjugate (helper ubiquitin, H), which cannot be part of the same ubiquitin chain as target ubiquitin^8^. Helper ubiquitin is proposed to bind a ubiquitin receptor on the proteasome, thus driving complex assembly through avidity, similarly to the UBL domain of Ubp6. Thus, even in the primed complex, productive docking of target ubiquitin to Ubp6 remains highly constrained by the requirement for helper ubiquitin, and will not take place for many proteasome substrates.

After docking of target ubiquitin (Extended Data Fig. 10, step 4), the final complex has the UBL of Ubp6 on Rpn1, target ubiquitin on Ubp6, and helper ubiquitin (or ubiquitin chain) engaged with a ubiquitin receptor. With the completion of this step, the proteasome assumes the si state and substrate degradation is suspended until deubiquitination takes place and target ubiquitin is released from the active site of Ubp6. After this release, the proteasome substrate has alternative fates; its degradation by the proteasome may proceed, or it may dissociate from the proteasome. The partitioning between these fates will likely depend on the number and arrangement of remaining ubiquitin groups on the proteasome substrate^8^. If, after deubiquitination, the substrate still carries multiple ubiquitin modifications, it may be subjected to successive rounds of deubiquitination by Ubp6.

Noncatalytic inhibition of the proteasome by Ubp6 is coupled to deubiquitination in that it is associated with the si state of the proteasome, in which ubiquitin is docked at the Ubp6 active site. The highly specific interactions that underlie proteasome inhibition by Ubp6 are exemplified by the ILR mutant Ubp6-AA and the Rpt1 activation loop mutant Rpt1-RK, both of which dramatically impair the noncatalytic effect even when ubiquitin has been chemically engineered to dock into the Ubp6 active site. Ubiquitin would normally not load onto Ubp6 in these mutants because they are defective in Ubp6 activation. The control of proteasome conformation by Ubp6 is expected to coordinate Ubp6 activity with other substrate processing events carried out by the proteasome. This mechanism grants time for Ubp6 to catalyze deubiquitination prior to substrate degradation, which should enhance the efficiency of chain removal by Ubp6.

#### Extended data figure legends

**Extended Data Fig. 1.**
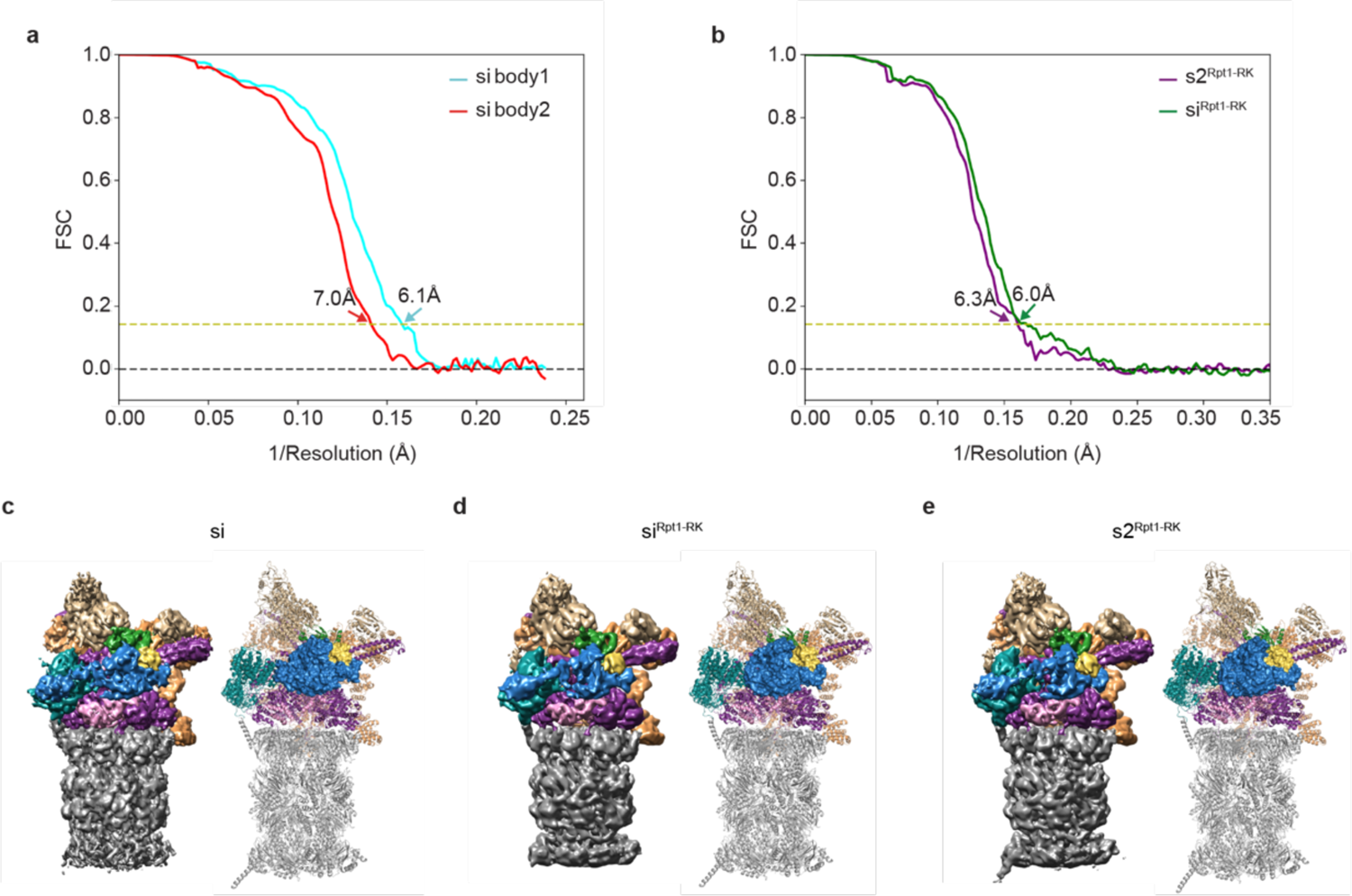
Cryo-EM reconstructions and fitted structural models of 26S proteasome-Ubp6 complexes. **a-b**, Fourier shell correlation curves of cryo-EM models in this study. **a**, Resolution of the reconstructions of si proteasomes, body1 (corresponding to the lid and CP) and body2 (corresponding to ATPase, Ubp6 and UbVS). **b**, Resolution of the reconstructions of s2^Rpt1-RK^ and si^Rpt1-RK^ proteasomes. Resolutions are determined on the basis of the gold standard Fourier shell correlation criterion (FSC=0.143, dotted yellow line). See Supplementary Table 1 for additional details. **c-e**, Overview of the cryo-EM reconstructions and the fitted structural model for the si state (**c**, EMDB-ID XXX, PDB-ID XXX, resolution 7.0 Å), the si^Rpt1-RK^ state (**d**, EMDB-ID XXX, PDB-ID XXX, 6.0 Å) and s2^Rpt1-RK^ (**e**, EMDB-ID XXX, PDB-ID XXX, 6.2 Å). The 26S proteasome is colored according to its subunits: Ubp6, blue; ubiquitin, yellow; Rpn11, green; Rpt1, pink; other Rpt subunits, purple; Rpn10 and other base subunits, tan; lid components, light brown; core particle, grey.

**Extended Data Fig. 2.**
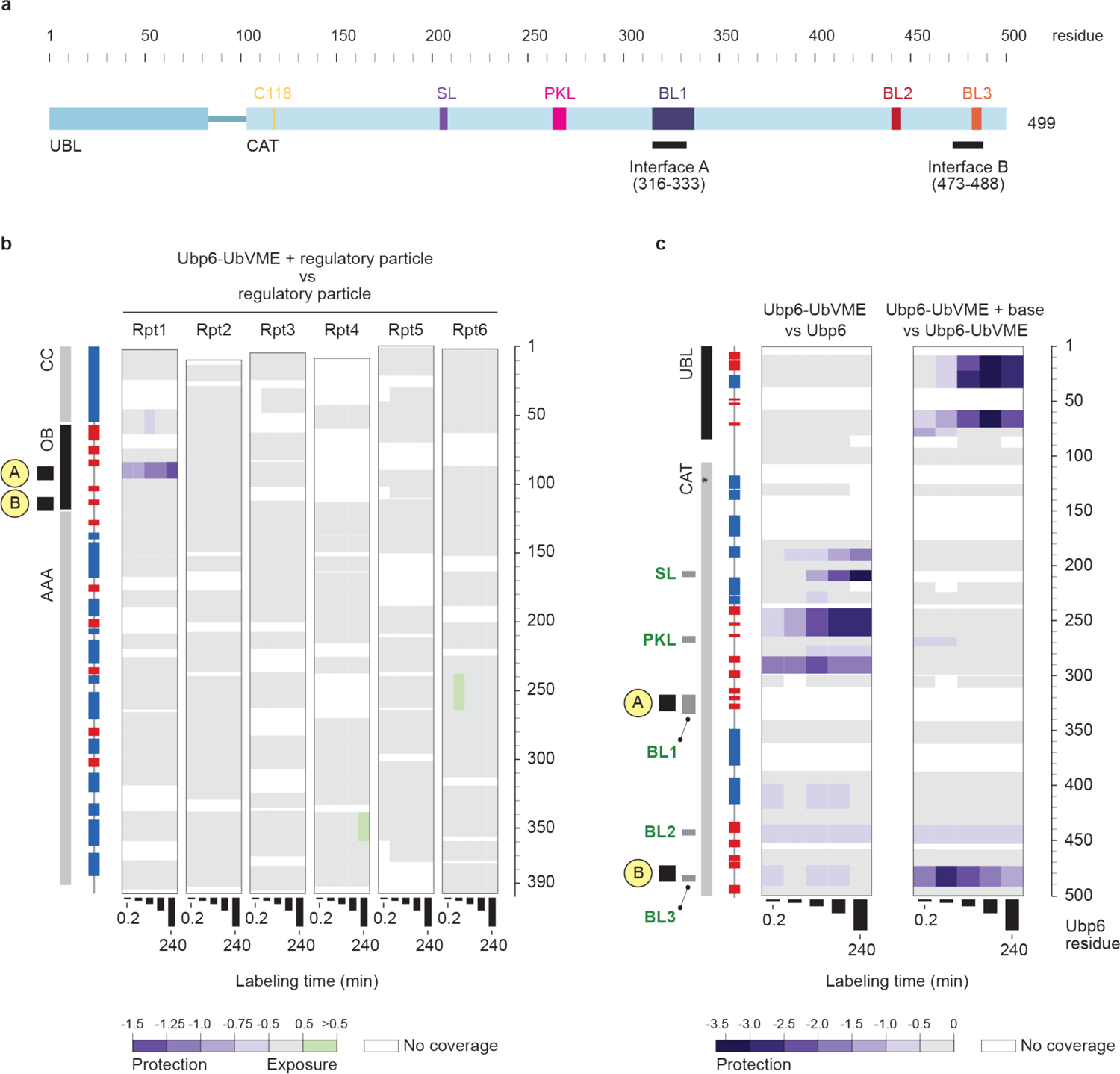
Hydrogen-deuterium exchange mass spectrometry confirms Ubp6 interaction with the OB domain of Rpt1. **a**, Schematic of functional elements of Ubp6. Ubiquitin-like (UBL) and catalytic (CAT) domains of Ubp6 are represented as blue boxes connected by a linker. Catalytic cysteine C118 and the five key loops are color-coded as in Fig. 1d. Amino acid residue numbers of Ubp6 are shown at top. Ubp6-Rpt1 interfaces are indicated as black bars at bottom. **b**, HXMS was performed to localize the Ubp6-Rpt1 interaction. This panel provides the complete data set from which Fig. 1c was abstracted. Deuterium exchange was monitored over four hours at time points of 0.17, 1, 10, 60, 240 min (represented by vertical bars). Deuteration differences of purified *ubp6Δ* regulatory particle (RP) alone versus RP in the presence of 5-fold molar excess of Ubp6-UbVME adduct were measured. Deuterium differences (D_RP+Ubp6-UbVME_ – D_RP alone_) are color-coded according to the scale at bottom, with the strongest shade of purple being most protected (i.e., the least amount of deuterium exchange) in the presence of Ubp6-UbVME. Domain organization at left and residue numbers at right refer to information determined from a master alignment between archaeal Rpt homolog proteasome-activating nucleotidase (PAN) and the six Rpt subunits in *S. cerevisiae*. α-helices and β-strands are depicted by blue and red boxes respectively. Interfaces A and B are shown as black boxes. Major protection is seen in the OB domain of Rpt1. This protection is unique to Rpt1; no other subunit (Rpt2-Rpt6) of the ATPase ring shows protection. **c**, Deuterium exchange in Ubp6. Domain organization of Ubp6, recovered from PDB:1WGG for the UBL and from the si structure for CAT, is shown at left, while residue numbers of Ubp6 are shown at right. All key Ubp6 loops are shown in grey boxes.The catalytic residue C118 is indicated by an asterisk. Differences in peptide deuteration over time are color-coded according to the scale at bottom. The left panel shows the differences (D_Ubp6-UbVME_ – D_Ubp6_) in deuteration of recombinant Ubp6-UbVME adduct versus Ubp6 alone. UbVME-induced protection is localized to a region of the catalytic domain. The right panel shows the differences (D_Ubp6-UbVME+base_ – D_Ubp6-UbVME alone_) in deuteration of the recombinant Ubp6-UbVME adduct alone versus in the presence of a 1.6-fold excess of recombinant proteasome base. Differences at each time point are color-coded according to the scale at the bottom. Strong protection within the UBL domain appears to represent its interaction with proteasomal subunit Rpn1^11^. For the CAT domain, only Interface B was shown to be protected; whereas peptides covering Interface A, which contains residues I329 and L330, were not resolved in this analysis.

**Extended Data Fig. 3.**
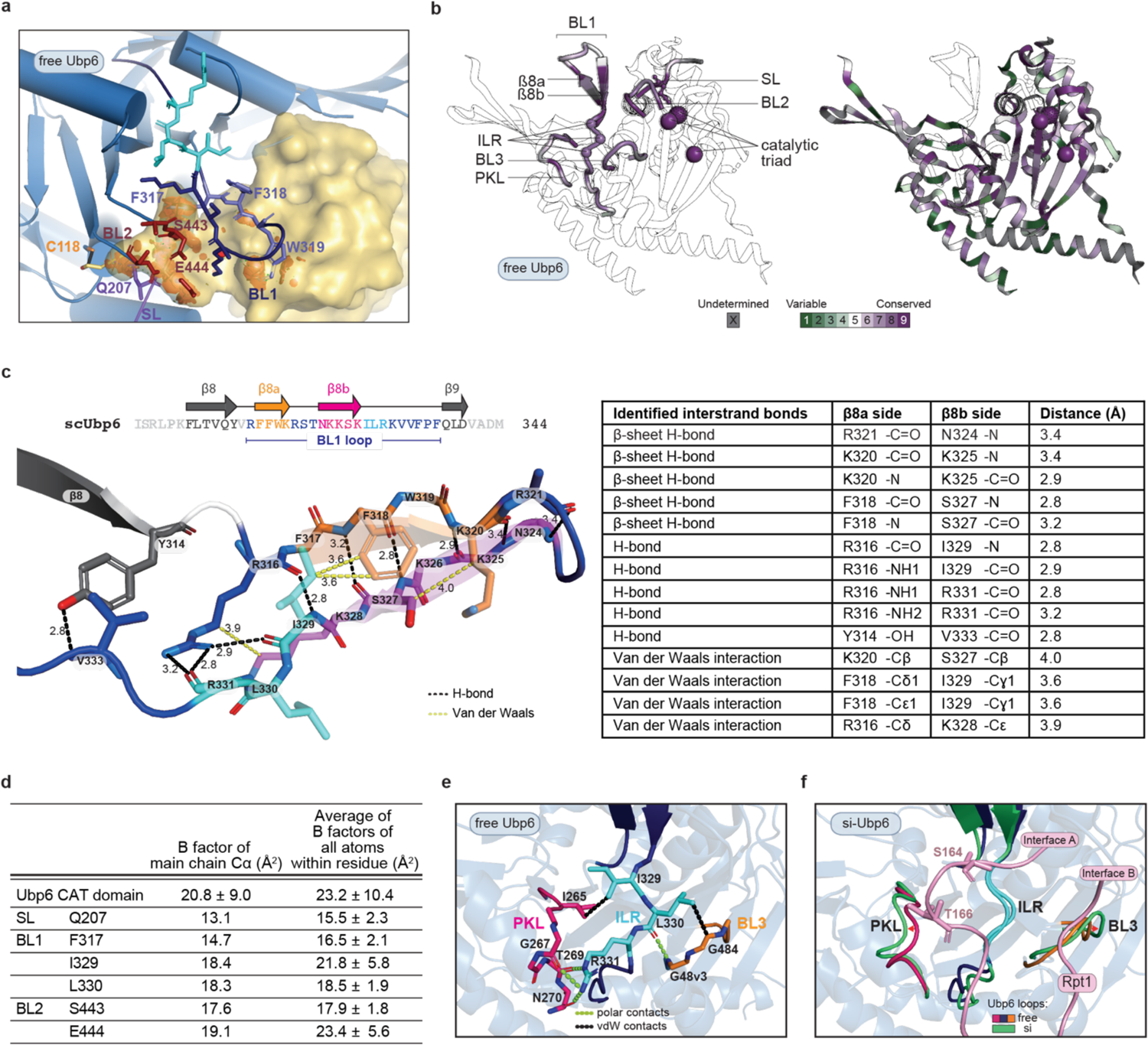
The blocking loop network of Ubp6. **a**, Zoomed-in view of ubiquitin-blocking loop clashes. The ubiquitin moiety from USP14-UbAl^13^ is superposed onto the free Ubp6 crystal structure. Steric hindrance, shown as embedded orange discs, can be seen between residues F317-W319 of BL1 (slate blue), BL2 (raspberry), Q207 of SL (purple) and ubiquitin (yellow). Specifically, prominent clashes from side chains of F317 and W319 and main chain of F318 in BL1 are evident, as are main chain and side chain clashes from S443 and E444 of BL2. The ILR activation element, which is within BL1, is displayed in cyan. In all panels, the isopeptide bond between the active site cysteine (orange) and C-terminus of ubiquitin (red) is represented in black. **b**, The blocking loop network of Ubp6 is evolutionarily conserved. Multiple sequence alignment of Ubp6 was performed with 50 orthologs from a diverse set of eukaryotes. Conservation analysis was then performed using the ConSurf server with Bayesian algorithm (https://consurf.tau.ac.il), where the output data represents the estimated evolutionary rate as graded conversation scores. Grade 9 (purple) indicates residues that are most conserved, Grade 1 (green) the most variable. For some residues (grey), evolutionary rate cannot be confidently computed (undetermined), for example when an alignment position has fewer than six ungapped amino acids or when a specific alignment interval contains conservation scores that span four or more grades. 91 out of 499 residues fell into the undetermined category. The nine color conservation scores were projected onto the crystal structure of free Ubp6 for visualization (PDB: 1VJV). (left) Evolutionary conservation of the three blocking loops and of BL3 and PKL. Residue Q207, the key ubiquitin-blocking residue of SL, is shown in ball and stick. The catalytic triad (C118, H447, N465) is rendered as spheres with the active site cysteine stippled. The remaining parts of Ubp6 are only outlined. (right) Overall conservation of the catalytic domain other than residues highlighted in panel *a*. Strong conservation is evident near the catalytic triad. **c**, Interactions within and around blocking loop 1 of Ubp6 stabilize the β-hairpin. Zoomed-in view of BL1 in free Ubp6 (PDB: 1VJV). Proposed main chain and side chain contacts between and adjacent to β-strands 8a (F317-K320, orange) and 8b (K324-K328, magenta) are indicated. Nitrogen and oxygen atoms are highlighted in skyblue and in red respectively. Two residues from the ILR element, I329 and L330, are labelled in cyan. Note that the stabilizing interstrand network extends from the ß-hairpin into and beyond the ILR element. A summary of interstrand interactions among residues Y314-V333 in free Ubp6 is given at right. **d**, Crystallographic B factors for free Ubp6 are consistent with minimal conformational dynamics in the SL loop, BL2, the ILR, and much of BL1 (excepting the distal tip of the hairpin). Apart from the ILR, residues shown in the table are those assigned as key blocking residues. For each residue specified, the B factors of main chain alpha carbon and the average of B factors of all atoms within the residue are given (PDB: 1VJV). B factor values less than 30 signify a low level of atomic fluctuation within the crystal. The top row gives averaged B factor values for the entire catalytic domain of Ubp6. **e**, Detail of the activation region at the BL1 base. In free Ubp6, the ILR is braced and held in place through multiple contacts with the adjacent PKL and BL3 loops. **f**, Comparison of the PKL-ILR-BL3 support in free Ubp6 (PDB: 1VJV) and complex si. Structures were aligned on the ILR (cyan). Loops from free Ubp6 were superimposed on si-Ubp6.

**Extended Data Fig. 4.**
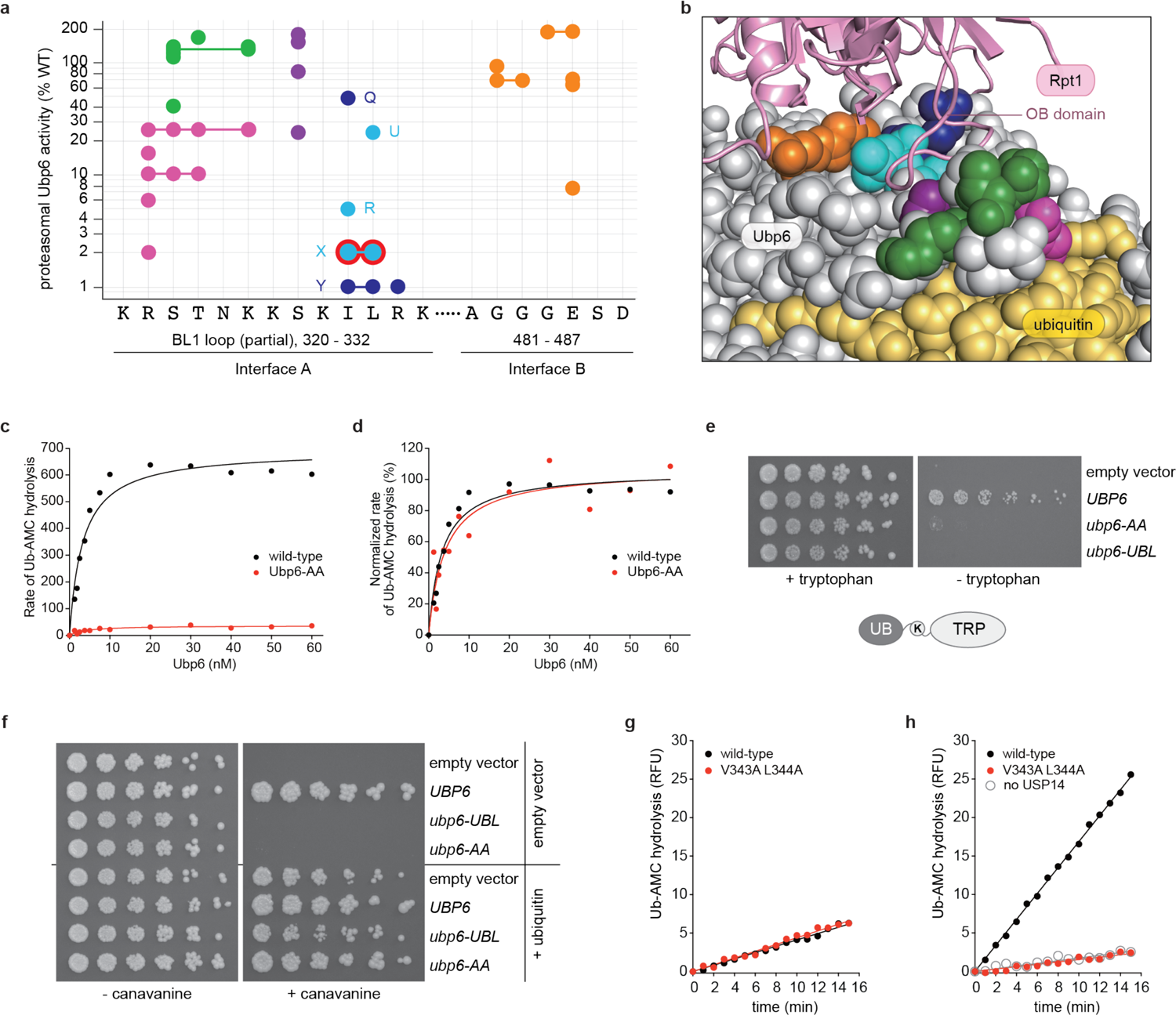
Characterization of *ubp6-I329A L330A* and *USP14-V343 L344* mutants. **a**, Proteasome-dependent activity of Ubp6 mutants mapped to the sequences of Interfaces A and B. Mutants color-coded as in Fig. 2a. **b**, Modeled Ubp6-Rpt1 interface with substituted residues color-coded as in Fig. 2a (PDB: this study). **c**, Concentration-dependence of Ubp6 activity on Ub-AMC (1 μM) in the presence of *ubp6Δ hul5Δ* proteasome (1 nM). The data were fit to a hyperbolic curve. **d**, Plot of the same data as at left, but with the calculated activity of the Ubp6-AA mutant normalized to that of wild-type. The calculated activity level at 60 nM was taken as the benchmark for normalization. The plot illustrates that the affinity of Ubp6 for the proteasome is only minimally altered by the mutation (apparent K_d_ of 3.3 nM for wild-type Ubp6 and 4.0 nM for Ubp6-AA), despite the strong defect in catalytic activity. The data were fit by nonlinear regression. The Ubp6-AA data are noisier than those of wild-type due to the low activity of the mutant enzyme. **e**, UB-K-TRP reporter stabilization by the *ubp6-AA* mutant. The UB-K-TRP fusion protein is co-translationally deubiquitinated, resulting in the expression of an unstable C-terminal fragment containing the Trp1 protein with lysine at its endoproteolytically processed N-terminus (K-TRP). K-TRP is efficiently degraded by the proteasome and thus has a short half-life, resulting in growth failure of the *ubp6Δ* strain in the absence of tryptophan. WT Ubp6 can restore growth on media lacking tryptophan by rescuing the reporter from degradation. In this assay, *ubp6Δ* mutant carrying an integrated UB-K-TRP reporter was transformed with plasmids expressing Ubp6 variants as indicated. Yeast cells were serially diluted, plated on media containing or lacking tryptophan, and incubated at 30°C for 3-7 days. **f**, Canavanine sensitivity test. Canavanine is an arginine analog that can be incorporated into newly synthesized proteins, thus impairing protein folding. *ubp6Δ* mutants are sensitive to canavanine, a phenotype that can result from the inability of cells to handle the increased burden of misfolded protein on proteasomes. In this assay, *ubp6Δ* mutants were transformed with plasmids expressing the indicated variants of *UBP6*. Transformants were serially diluted and transferred to agar plates containing 100 μM CuSO_4_ in the presence or absence of canavanine at 1.5 μg/mL. Plates were incubated at 30°C for 3-7 days. The canavanine-sensitivity of *ubp6* mutants can be suppressed by ubiquitin overexpression: ubiquitin expression is driven by the *CUP1* promoter and induced by supplemented CuSO_4_. *UBP6*, wild-type UBP6; *UBL,* UBL domain of Ubp6. **g**-**h**, The data shown here were used to generate Fig. 2e. **g**, Basal USP14 activity. Ub-AMC hydrolysis assay of wild-type USP14 (1 μM) in its free form, together with the V343A L344A double mutant. **h**, Proteasomal activation of deubiquitination by USP14. Ub-AMC hydrolysis assay of WT and mutant USP14 (8 nM) in the presence of human proteasome (1 nM) pretreated with ubiquitin vinyl sulfone (UbVS) to eliminate UCH-L5 activity. “No USP14,” UbVS-treated human proteasome alone.

**Extended Data Fig. 5.**
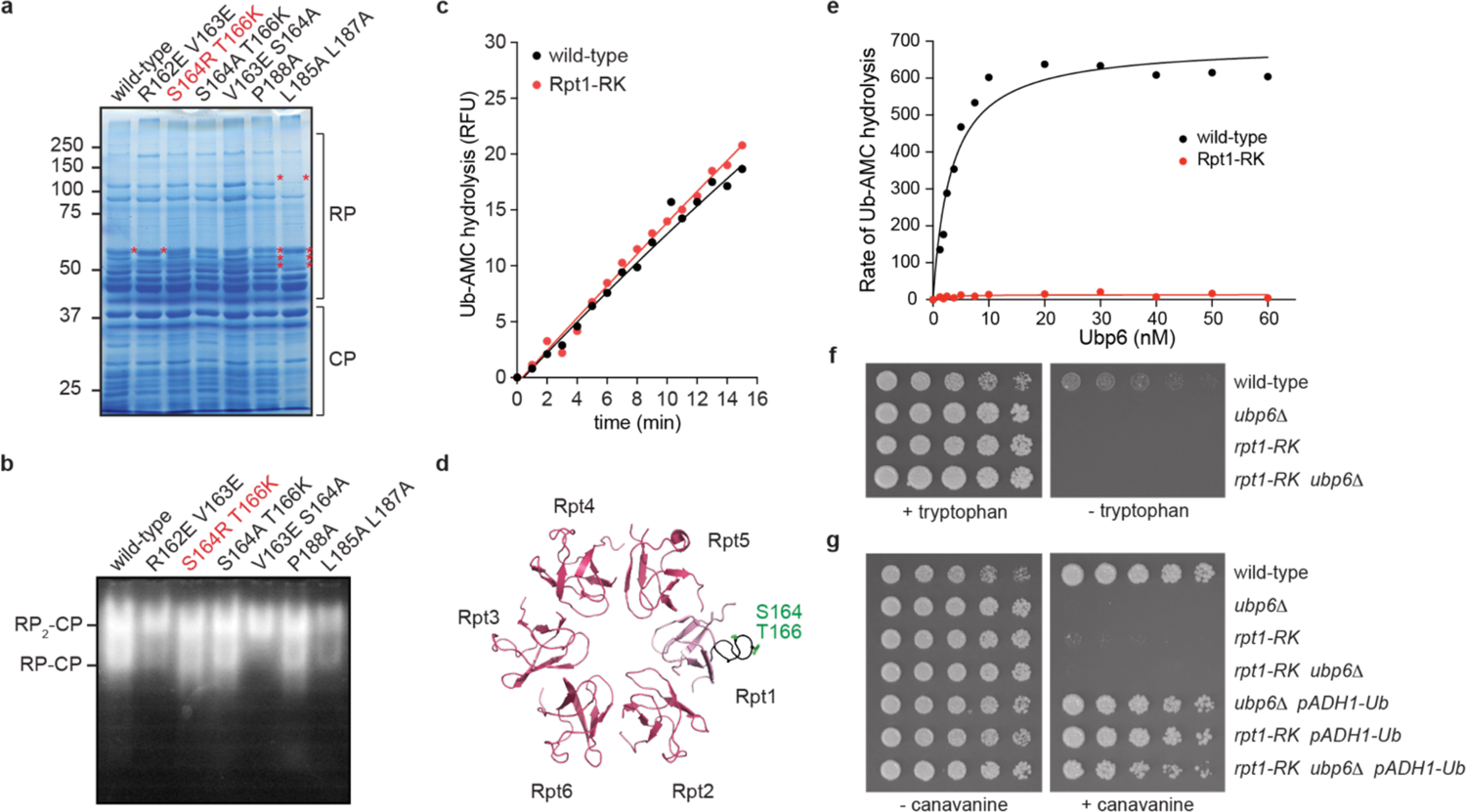
An Rpt1 mutant defective in Ubp6 activation. **a**, Rpt1-RK mutant proteasomes (S164R T166K) do not exhibit assembly defects. Coomassie-blue-stained SDS-PAGE gel showing the subunit composition of purified proteasomes from yeast strains bearing the indicated *rpt1* mutations. The mutant proteasome of interest, Rpt1-RK, shows an electrophoretic profile that is indistinguishable from wild-type. Protein bands reflecting altered subunit composition in other mutants are marked by asterisks. **b**, Purified proteasome complexes were resolved by 3.5% native PAGE, and active species were visualized by an in-gel Suc-LLVY-AMC assay. RP_2_-CP and RP-CP are doubly and singly-capped proteasomes, respectively. Proteasome from *rpt1-RK* mutant shows a profile comparable to that from wild-type. Proteasome assembly is perturbed in samples from *rpt1-R162E-V163E*, *rpt-V163E-S164A*, and *rpt1-L185A-L187A* mutant strains. **c**, Time course showing the basal Ub-AMC hydrolytic activity of proteasomes purified from *ubp6Δ hul5Δ* (nominally wild-type) or *rpt1-RK ubp6Δ hul5Δ* strains. 1 μM Ub-AMC was used in this assay. Ubp6-independent “background” hydrolytic activity shown is comparable between the two proteasome samples. **d**, OB domain, top view. The L34 activation loop of Rpt1 is in black, S164 and T166 in green. (PDB: 6EF3). **e**, The effect of Ubp6 concentration on Ub-AMC hydrolysis in the presence of proteasome (1 nM) purified from *ubp6Δ hul5Δ* (nominally wild-type) or *rpt1-RK ubp6Δ hul5Δ* strain. Data were fit to a hyperbolic curve, yielding an apparent *K*_d_ value of ∼3.3 nM and ∼2.0 nM for wild-type and Rpt1-RK proteasome, respectively. **f**, UB-K-TRP reporter stabilization by *rpt1-RK* mutation. Yeast cells were serially diluted and plated on media containing or lacking tryptophan. **g**, Wild-type, *ubp6Δ*, and *rpt1-RK* mutants were serially diluted and spotted onto agar plates in the presence or absence of canavanine (1.5 μg/mL). The *pADH1-Ub* transgene is integrated into the *UBP6* locus and expresses ubiquitin from the *ADH1* promoter to compensate for the ubiquitin deficiency of the *ubp6* null.

**Extended Data Fig. 6.**
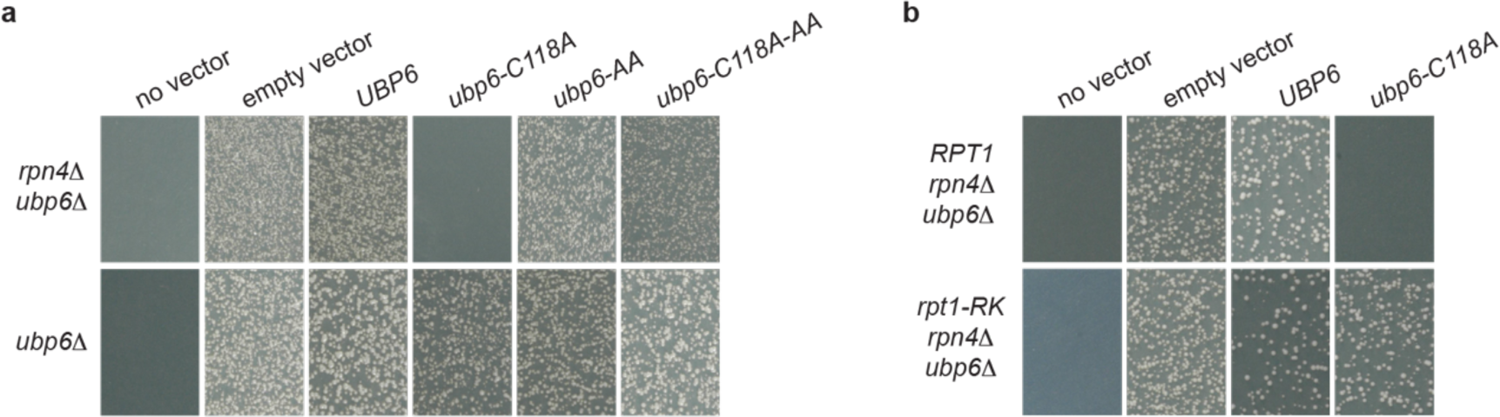
*ubp6-AA* and *rpt1-RK* mutants are impaired *in vivo* in the Ubp6 noncatalytic effect. **a**, *In vivo* assay of the Ubp6 noncatalytic effect^48^. Plasmids expressing variants of Ubp6 were transformed into either *ubp6Δ rpn4Δ* or *ubp6Δ* yeast strains, and colony formation was recorded after incubation at 30°C for 3-4 days. Loss of noncatalytic activity was seen in *ubp6-C118A-AA*. **b**, The noncatalytic effect of *ubp6-C118A* is abrogated by the *rpt1-RK* mutation.

**Extended Data Fig. 7.**
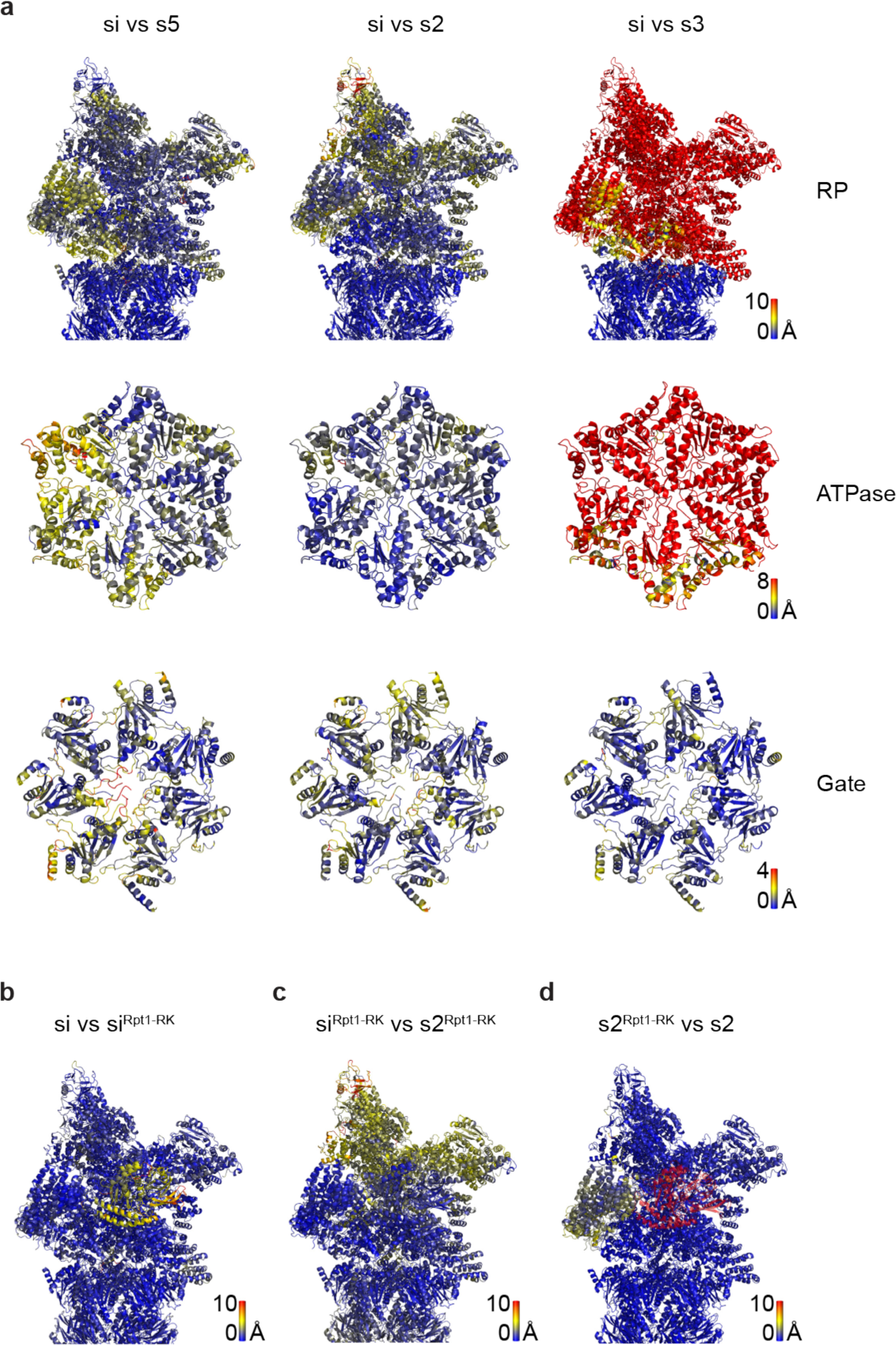
Structural comparisons of proteasome conformational states observed in the presence and absence of Ubp6. All comparisons shown are visualized by coloring scheme based on the residuewise root mean square deviation (RMSD). States that are compared are aligned with respect to the CP. The RMSD of each atom is calculated and the RMSD values of all atoms in one residue are averaged. Then the residues are colored based on the spectral distribution of averaged residuewise RMSD. Residues with small RMSD suggest high similarity between compared structures and are shown in blue, while those with large RMSD, suggesting low similarity between compared structures, are shown in red. **a**, Structural differences between si and s5, s2, or s3 proteasomes. Shown here are the si proteasomes, which is used as the reference for comparison. si-state proteasomes combine structural elements of previously reported proteasomal states without bound Ubp6. The RP of si is most similar to s5 (PDB: 6FVX), the ATPase to s2 (PDB: 6FVU), and the gate to s3 (PDB: 6FVV)^17^. **b**, Structural differences between si and si^Rpt1-RK^. Shown is si. The Rpt1-RK mutations evoke substantial differences only in Ubp6 positioning; for si state proteasomes, the overall proteasomal structure remains unaffected. **c**, Structural differences between si^Rpt1-RK^ and s2^Rpt1-RK^. Shown is si^Rpt1-RK^. The two identified states of the Rpt1-RK mutant are almost identical to each other with respect to Ubp6 positioning. **d**, Structural differences between the Ubp6-containing s2^Rpt1-RK^ and the Ubp6-free s2 (PDB: 6FVU)^17^. Shown is s2^Rpt1-RK^. Ubp6 binding per se does not evoke substantial structural differences for this mutant. The slight differences indicated by the yellow coloring of Rpn1 are due to Rpn1’s lower local resolution.

**Extended Data Fig. 8.**
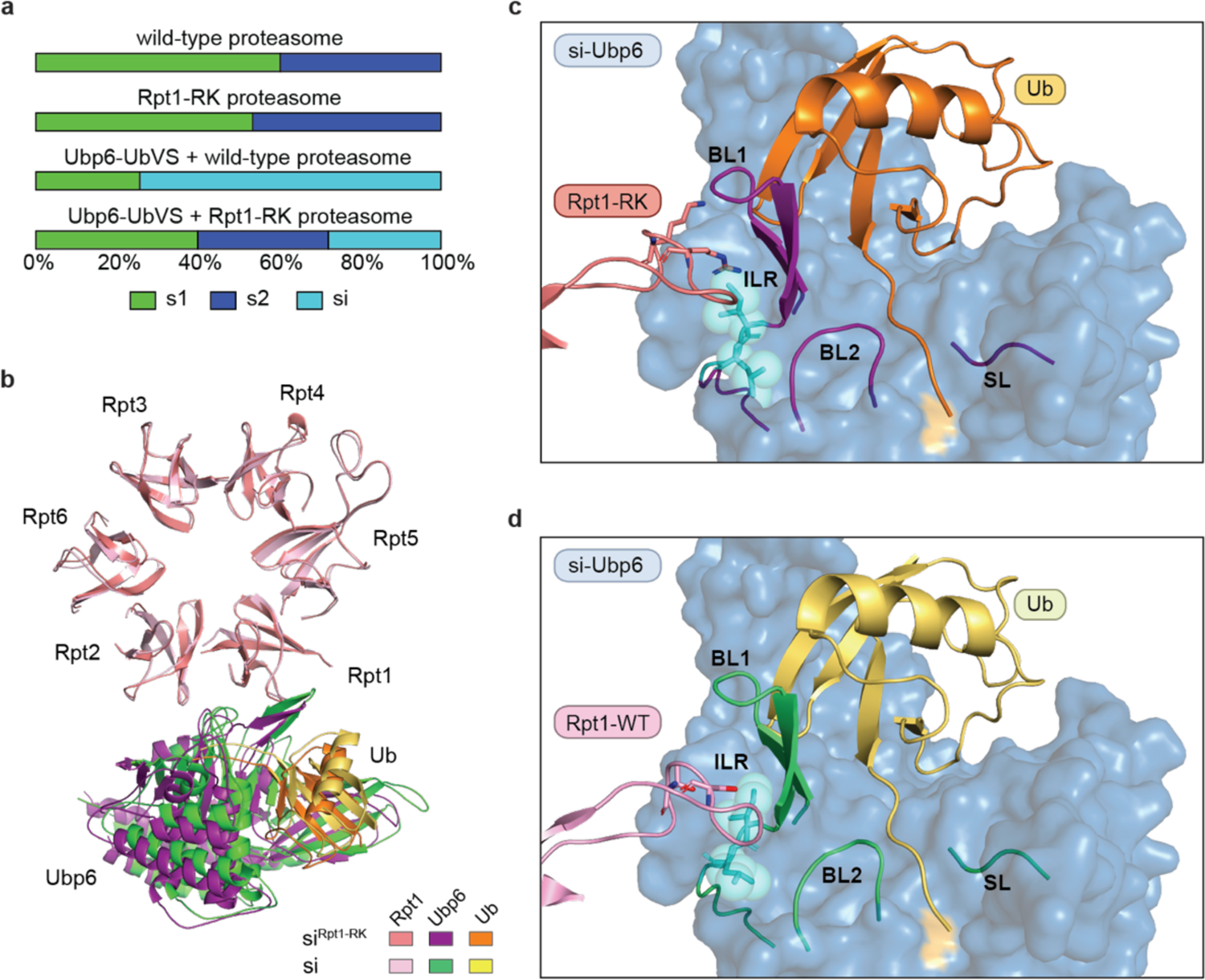
The *Rpt1-S164R T166K* double mutation blocks Ubp6 activation and shifts the ternary complex from the inhibited si-state to the s2 state. a, Effect of the *rpt1-RK* mutation on the proteasome state distribution. b, The *Rpt1-RK* mutation evokes a repositioning of Ubp6-UbVS relative to the proteasome. The si and si^Rpt1-RK^ complexes were aligned by their OB rings. c, Zoomed-in view of the contact interface between the activation loop of Rpt1-RK (salmon) and BL1 (purple) of Ubp6 in the si^Rpt1-RK^ proteasome. The positively charged bulky side chains of residues R164 and K166 on the Rpt1 mutant result in less extensive contacts between the activation loop and the ILR. d, Zoomed-in view of the contact interface between the activation loop of wild-type Rpt1 (pink) and the ILR element (cyan) within BL1 of Ubp6.

**Extended Data Fig. 9.**
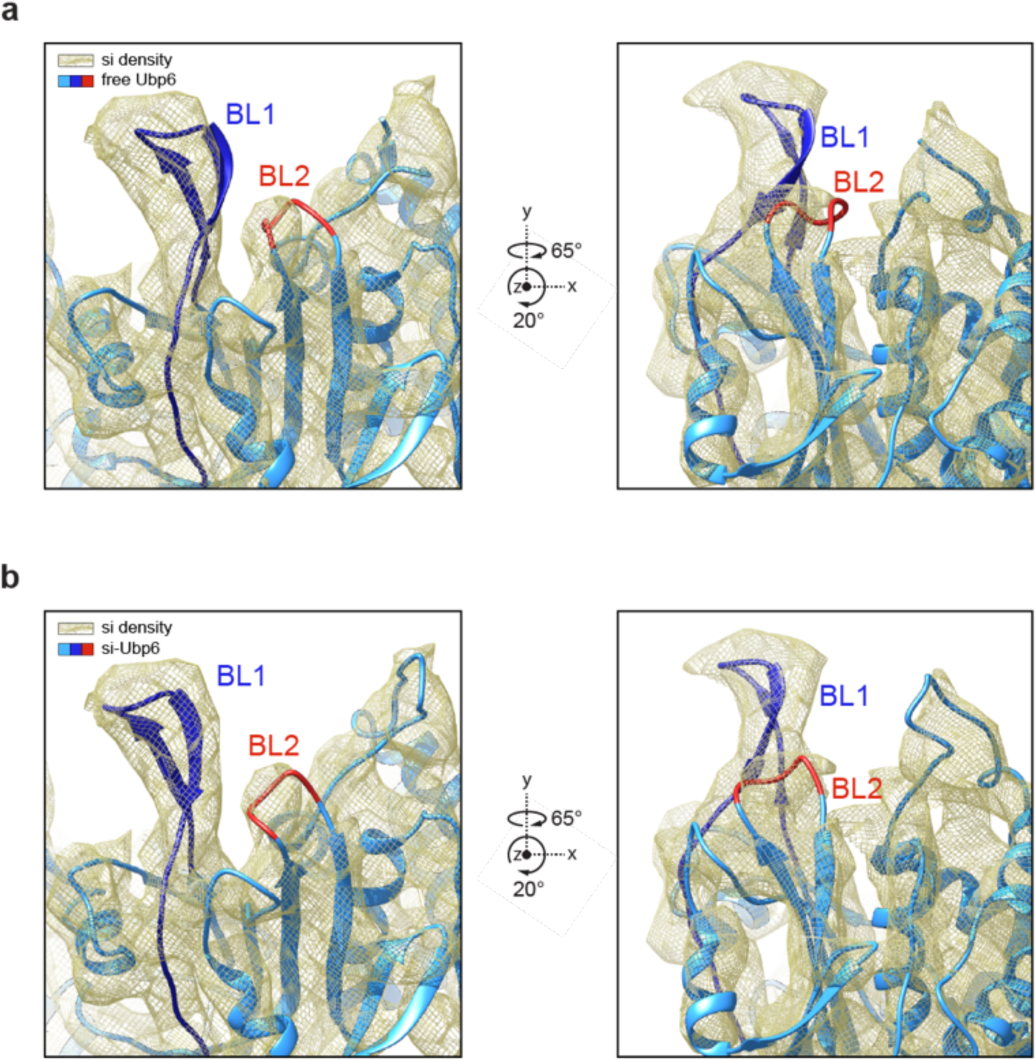
Detailed view of the conformational change of BL1 and BL2 loops. **a**, Cryo-EM density map of the si state proteasome is shown with free Ubp6 from the X-ray crystal structure (PDB: 1VJV). **b**, Corresponding model of Ubp6 in si proteasomes. Ubp6 structures excluding BL1(316-337), BL2(441-445) and peripheral two helices (345-416) were fitted into the si density. BL1 and BL2 are highlighted in blue and red respectively. The BL1 and BL2 loops of free Ubp6 do not fit well with the Ubp6 Cryo-EM density from the si proteasome.

**Extended Data Fig. 10.**
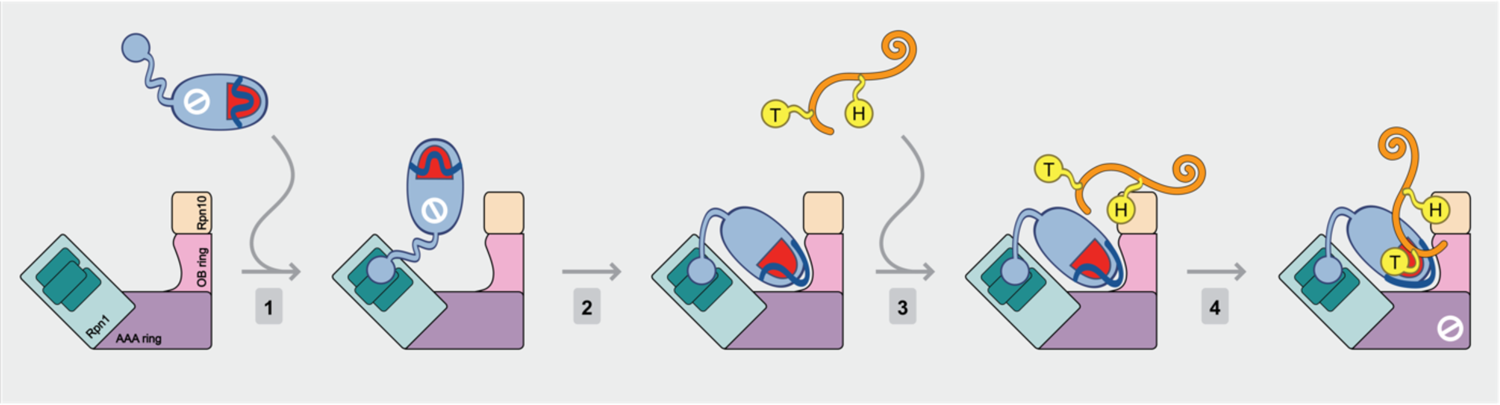
Model for assembly of the Ubp6 catalytic complex. Free Ubp6 has an inactive conformation (slashed circle), its active site (red semioval) being blocked by BL1, BL2, and SL (blue wave). Step 1, Ubp6 and the proteasome are complexed via the Ubp6^UBL^-Rpn1 interaction^2^. For clarity, only a fraction of proteasome subunits is represented. The Ubp6^UBL^-Rpn1 interaction does not activate Ubp6 (ref. ^10^) but is proposed to promote association of the Ubp6 catalytic domain to Rpt1 through avidity (step 2). Ubp6-Rpt1 interaction in this context may partially destabilize the blocking loop network to enable ubiquitin loading; this remains conjectural. Activated Ubp6 remains highly selective in that it will efficiently cleave only ubiquitin-protein conjugates that carry more than one ubiquitin modification^8^. Thus, a second docking event wherein substrate-bound “helper ubiquitin” (H) is docked at a ubiquitin receptor such as Rpn10, is required (step 3). With step 4, docking of the “target ubiquitin” (T), the catalytic complex is assembled: blocking loops are displaced more completely, and the proteasome assumes the si state, imposing noncatalytic proteasome inhibition. Thus, assembly of a competent catalytic complex essentially requires three docking events involving ubiquitin or ubiquitin-like protein domains: Ubp6^UBL^ at Rpn1, helper ubiquitin at a ubiquitin receptor, and target ubiquitin in the Ubp6 active site. For additional comments, please see the *Supplementary Discussion*.

**Supplementary Table 1.** Cryo-EM data collection, refinement and validation statistics

|  | si (body1)<br>(EMDB-xxxx)<br>(PDB xxxx) | si (body2)<br>(EMDB-xxxx)<br>(PDB xxxx) | si<sup>Rpt1-RK</sup> | s2<sup>Rpt1-RK</sup> |
| --- | --- | --- | --- | --- |
| **Data collection and processing** |  |  |  |  |
| Magnification | 22500 | 22500 | 18000 | 18000 |
| Voltage (kV) | 300 | 300 | 300 | 300 |
| Electron exposure (e-/Å<sup>2</sup>) | 60 | 60 | 35 | 35 |
| Defocus range (μm) | -1.0 to -2.5 | -1.0 to -2.5 | -1.8 to -3.0 | -1.8 to -3.0 |
| Pixel size (Å) | 2.10 | 2.10 | 1.38 | 1.38 |
| Symmetry imposed | C1 | C1 | C1 | C1 |
| Initial particle images (no.) | 112802 | 112802 | 231296 | 231296 |
| Final particle images (no.) | 88243 | 88243 | 64766 | 74842 |
| Map resolution (Å) | 6.1 | 7 | 6 | 6.3 |
| FSC threshold | 0.143 | 0.143 | 0.143 | 0.143 |
| Map resolution range (Å) | 4.3-12 | 6-20.8 | 4.8– 17.8 | 4.6 – 15 |
| **Refinement** |  |  |  |  |
| Initial model used (PDB code) | 6FVX (Lid)<br>6FVV (CP) | 6FVU (ATPase)<br>1VJV (Ubp6)<br>2AYO (UbVS) | si | 6FVU<br>1VJV (Ubp6)<br>2AYO (UbVS) |
| Model resolution (Å) | 7.1 | 8.1 | 7.5 | 8.6 |
| FSC threshold | 0.5 | 0.5 | 0.5 | 0.5 |
| Map sharpening *B* factor (Å<sup>2</sup>) | -306 | -520 | -263 | -304 |
| Model composition |  |  |  |  |
| Non-hydrogen atoms | 90,304 | 22,546 | 112,857 | 112,954 |
| Protein residues | 11,480 | 2,831 | 14,311 | 14,323 |
| Ligands | 0 | 12 | 12 | 12 |
| R.m.s. deviations |  |  |  |  |
| Bond lengths (Å) | 0.014 | 0.011 | 0.035 | 0.008 |
| Bond angles (°) | 1.996 | 1.887 | 2.849 | 1.527 |
| Validation |  |  |  |  |
| MolProbity score | 2.33 | 2.15 | 2.52 | 2.57 |
| Clashscore | 11.62 | 7.09 | 8.78 | 11.02 |
| Poor rotamers (%) | 2.79 | 1.94 | 6.29 | 6.90 |
| Ramachandran plot |  |  |  |  |
| Favored (%) | 93.80 | 90.46 | 93.19 | 94.52 |
| Allowed (%) | 5.42 | 8.19 | 5.78 | 4.64 |
| Disallowed (%) | 0.78 | 1.35 | 1.03 | 0.84 |

**Supplementary Table 2.**
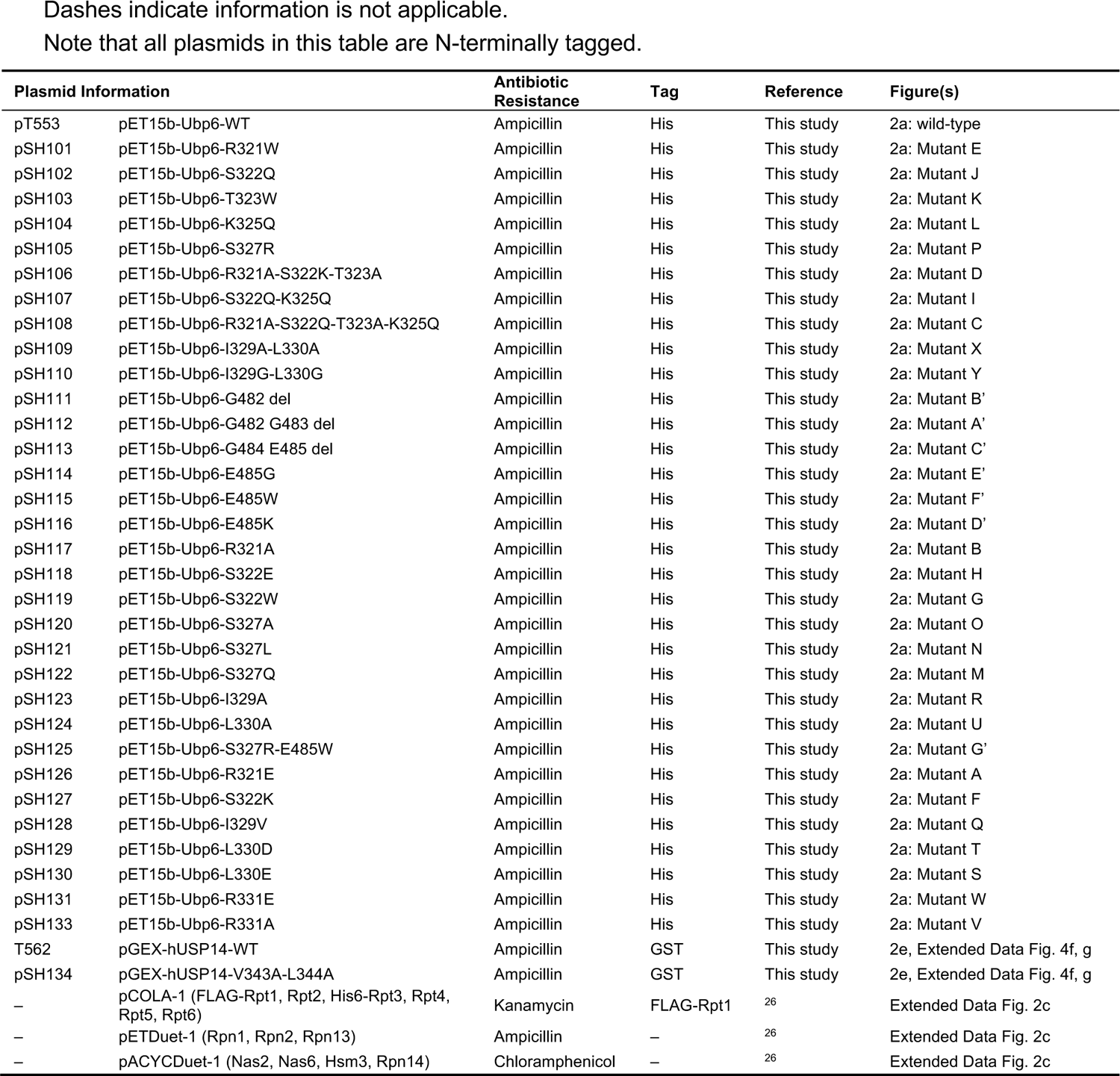
Plasmid information for recombinant protein expression.

| Plasmid Information |  | Antibiotic Resistance | Tag | Reference | Figure(s) |
| --- | --- | --- | --- | --- | --- |
| pT553 | pET15b-Ubp6-WT | Ampicillin | His | This study | 2a: wild-type |
| pSH101 | pET15b-Ubp6-R321W | Ampicillin | His | This study | 2a: Mutant E |
| pSH102 | pET15b-Ubp6-S322Q | Ampicillin | His | This study | 2a: Mutant J |
| pSH103 | pET15b-Ubp6-T323W | Ampicillin | His | This study | 2a: Mutant K |
| pSH104 | pET15b-Ubp6-K325Q | Ampicillin | His | This study | 2a: Mutant L |
| pSH105 | pET15b-Ubp6-S327R | Ampicillin | His | This study | 2a: Mutant P |
| pSH106 | pET15b-Ubp6-R321A-S322K-T323A | Ampicillin | His | This study | 2a: Mutant D |
| pSH107 | pET15b-Ubp6-S322Q-K325Q | Ampicillin | His | This study | 2a: Mutant I |
| pSH108 | pET15b-Ubp6-R321A-S322Q-T323A-K325Q | Ampicillin | His | This study | 2a: Mutant C |
| pSH109 | pET15b-Ubp6-I329A-L330A | Ampicillin | His | This study | 2a: Mutant X |
| pSH110 | pET15b-Ubp6-I329G-L330G | Ampicillin | His | This study | 2a: Mutant Y |
| pSH111 | pET15b-Ubp6-G482 del | Ampicillin | His | This study | 2a: Mutant B' |
| pSH112 | pET15b-Ubp6-G482 G483 del | Ampicillin | His | This study | 2a: Mutant A' |
| pSH113 | pET15b-Ubp6-G484 E485 del | Ampicillin | His | This study | 2a: Mutant C' |
| pSH114 | pET15b-Ubp6-E485G | Ampicillin | His | This study | 2a: Mutant E' |
| pSH115 | pET15b-Ubp6-E485W | Ampicillin | His | This study | 2a: Mutant F' |
| pSH116 | pET15b-Ubp6-E485K | Ampicillin | His | This study | 2a: Mutant D' |
| pSH117 | pET15b-Ubp6-R321A | Ampicillin | His | This study | 2a: Mutant B |
| pSH118 | pET15b-Ubp6-S322E | Ampicillin | His | This study | 2a: Mutant H |
| pSH119 | pET15b-Ubp6-S322W | Ampicillin | His | This study | 2a: Mutant G |
| pSH120 | pET15b-Ubp6-S327A | Ampicillin | His | This study | 2a: Mutant O |
| pSH121 | pET15b-Ubp6-S327L | Ampicillin | His | This study | 2a: Mutant N |
| pSH122 | pET15b-Ubp6-S327Q | Ampicillin | His | This study | 2a: Mutant M |
| pSH123 | pET15b-Ubp6-I329A | Ampicillin | His | This study | 2a: Mutant R |
| pSH124 | pET15b-Ubp6-L330A | Ampicillin | His | This study | 2a: Mutant U |
| pSH125 | pET15b-Ubp6-S327R-E485W | Ampicillin | His | This study | 2a: Mutant G' |
| pSH126 | pET15b-Ubp6-R321E | Ampicillin | His | This study | 2a: Mutant A |
| pSH127 | pET15b-Ubp6-S322K | Ampicillin | His | This study | 2a: Mutant F |
| pSH128 | pET15b-Ubp6-I329V | Ampicillin | His | This study | 2a: Mutant Q |
| pSH129 | pET15b-Ubp6-L330D | Ampicillin | His | This study | 2a: Mutant T |
| pSH130 | pET15b-Ubp6-L330E | Ampicillin | His | This study | 2a: Mutant S |
| pSH131 | pET15b-Ubp6-R331E | Ampicillin | His | This study | 2a: Mutant W |
| pSH133 | pET15b-Ubp6-R331A | Ampicillin | His | This study | 2a: Mutant V |
| T562 | pGEX-hUSP14-WT | Ampicillin | GST | This study | 2e, Extended Data Fig. 4f, g |
| pSH134 | pGEX-hUSP14-V343A-L344A | Ampicillin | GST | This study | 2e, Extended Data Fig. 4f, g |
| – | pCOLA-1 (FLAG-Rpt1, Rpt2, His6-Rpt3, Rpt4, Rpt5, Rpt6) | Kanamycin | FLAG-Rpt1 | <sup>26</sup> | Extended Data Fig. 2c |
| – | pETDuet-1 (Rpn1, Rpn2, Rpn13) | Ampicillin | – | <sup>26</sup> | Extended Data Fig. 2c |
| – | pACYCDuet-1 (Nas2, Nas6, Hsm3, Rpn14) | Chloramphenicol | – | <sup>26</sup> | Extended Data Fig. 2c |

**Supplementary Table 3.**
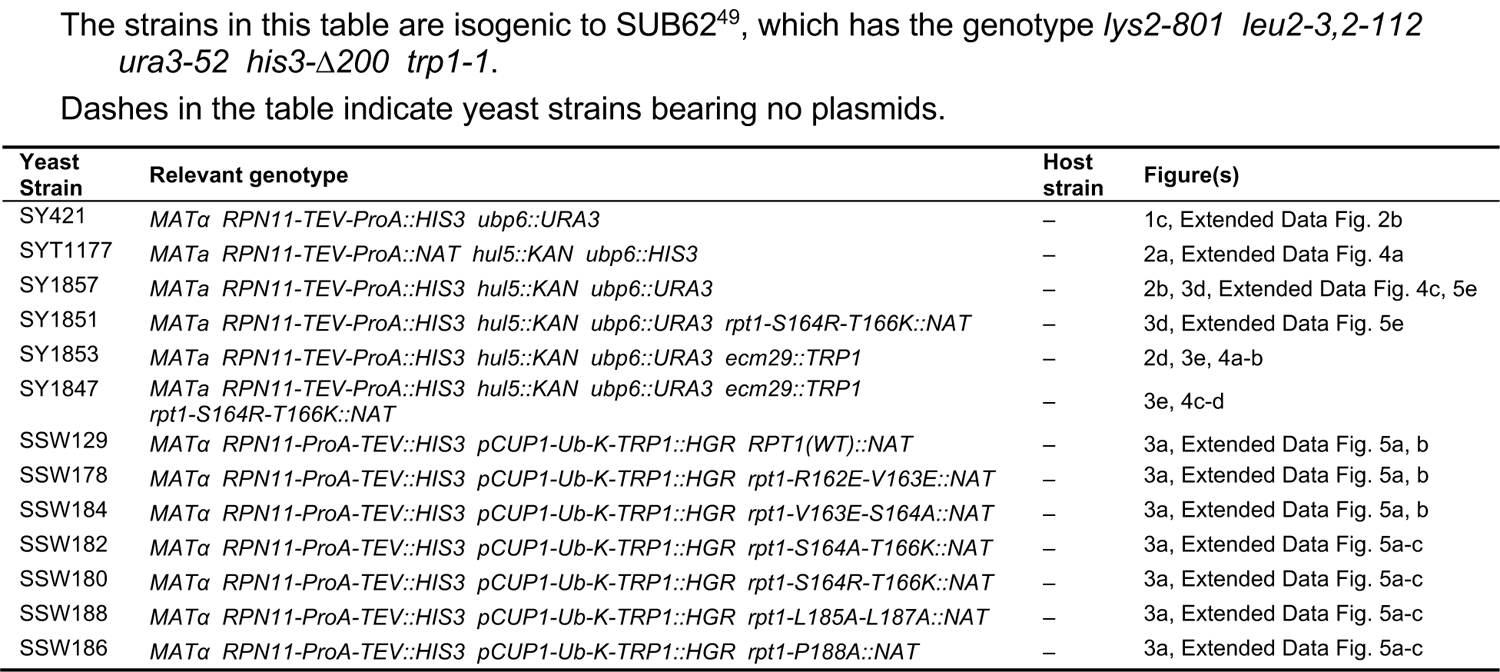
Yeast proteasome purifications strains used for biochemical assays.

**Supplementary Table 4.** Yeast proteasome purifications strains used for cryo-electron microscopy studies.

| Yeast Strain | Relevant genotype | Figure(s) |
| --- | --- | --- |
| YYS40<sup>50</sup> | *MATa RPN11-3xFLAG-HIS3* | 1, 2, 3, 5, Extended Data Fig. 1, 4, 7, 8, 9 |
| SSW234 | *MATa RPN11-3xFLAG-HIS3 rpt1-S164R-T166K::NAT* | Extended Data Fig. 1, 7, 8 |

**Supplementary Table 5.** Hydrogen deuterium exchange mass spectrometry data summary and list of experimental parameters.

| Data Set | Measure uptake in RP<br>(Fig. 1c; Extended Data Fig. 2b) | Measure uptake in Ubp6<br>(Extended Data Fig. 2c) |
| --- | --- | --- |
| States analyzed | 1. RP alone<br>2. RP + Ubp6UbVME (1:5) | 3. Ubp6 alone<br>4. Ubp6UbVME alone<br>5. Ubp6UbVME + base (1:2) |
| HDX reaction details <sup>a</sup> | Final D<sub>2</sub>O concentration = 92.2%, pH<sub>read</sub> = 7.15, 21 °C |  |
| HDX time course | 0.167, 1, 10, 60, 240 minutes |  |
| HDX controls | 3 undeuterated per state |  |
| Back-exchange | 30-35% |  |
| Filtering parameters | 0.3 products/amino acid;<br>2 consecutive products; 10 ppm error | 0.3 products/amino acid;<br>1 consecutive product; 10 ppm error |
| Number of peptides <sup>b</sup> | Rpt1: 45 followed; 62 identified<br>Rpt2: 42 followed; 54 identified<br>Rpt3: 42 followed; 68 identified<br>Rpt4: 32 followed; 42 identified<br>Rpt5: 48 followed; 62 identified<br>Rpt6: 53 followed; 63 identified | 71 followed; 87 identified |
| Sequence coverage <sup>b</sup> | Rpt1: 69.2%<br>Rpt2: 77.1%<br>Rpt3: 80.1%<br>Rpt4: 56.8%<br>Rpt5: 74.9%<br>Rpt6: 83.7% | 73.30% |
| Average peptide length/<br>Redundancy <sup>b</sup> | Rpt1: 15.7; 2.15<br>Rpt2: 16.4; 2.03<br>Rpt3: 14.4; 1.77<br>Rpt4: 15.4; 1.99<br>Rpt5: 13.7; 2.02<br>Rpt6: 16.0; 2.50 | 13.6; 2.53 |
| Replicates | 2 technical per state |  |
| Repeatability | +/- 0.15 relative Da |  |
| Meaningful Differences | > 0.5 Da |  |

<sup>a</sup> 12-fold dilution with labeling buffer (10 mM HEPES-NaOD [pD 7.5], 50 mM NaCl, 50 mM KCl, 5 mM MgCl<sub>2</sub>, 0.5 mM EDTA, 0.5 mM ATP, 1 mM DTT, 10% glycerol, 99.9% D<sub>2</sub>O). 1:1 dilution with quench buffer (0.8 M guanidinium chloride, 0.8% [v/v] formic acid, H<sub>2</sub>O, pH 2.0)
<sup>b</sup> Averages of separately processed replicates

**Supplementary Table 6.** Antibodies used in this study.

| Antibody |  | Working Dilution | Source | Reference |
| --- | --- | --- | --- | --- |
| HA epitope YPYDVPDYA | HRP-conjugated | 1:1,000 | Sigma | 12013819001 |
| Ubp6 | Rabbit polyclonal | 1:50,000 | Finley lab | <sup>11</sup> |
| Rpn13 | Rabbit polyclonal | 1:10,000 | Finley lab | This study |
| Rpn8 | Rabbit polyclonal | 1:15,000 | Finley lab | <sup>48</sup> |

**Supplementary Table 7.** Plasmid information for yeast transformation.

| Plasmid Information |  | Backbone | Reference | Figure(s) |
| --- | --- | --- | --- | --- |
| YCplac33 | CEN4/URA3, empty vector | n.a. | <sup>51</sup> | Extended Data Fig. 4, 6 |
| pJH80 | UBP6-WT | YPplac33 | <sup>15</sup> | Extended Data Fig. 4, 6 |
| pJH81 | ubp6-C118A | YPplac33 | <sup>15</sup> | Extended Data Fig. 6 |
| pySH01 | ubp6-I329A-L330A | YPplac33 | This study | Extended Data Fig. 4, 6 |
| pySH04 | ubp6-C118A-I329A-L330A | YPplac33 | This study | Extended Data Fig. 6 |
| pySH07 | ubp6-UBL | YPplac33 | This study | Extended Data Fig. 4 |
| YEp46Δ | 2μ/TRP1, empty vector | n.a. | <sup>52</sup> | Extended Data Fig. 4 |
| YEp96 | pCUP1-Ub | YEp46Δ | <sup>52</sup> | Extended Data Fig. 4 |

**Supplementary Table 8.** Yeast strains used for Ub-K-Trp assay.

| Yeast Strain | Relevant genotype | Host strain | Figure(s) |
| --- | --- | --- | --- |
| sJH138<sup>15</sup> | *MATa pCUP1-Ub-K-TRP1::NAT ubp6::KAN* | – | – |
| ySH29 | *MATa pCUP1-Ub-K-TRP1::NAT ubp6::KAN (YCplac33 [CEN4/URA3])* | sJH138 | Extended Data Fig. 4d |
| ySH30 | *MATa pCUP1-Ub-K-TRP1::NAT ubp6::KAN (pJH80: UBP6-WT [CEN4/URA3])* | sJH138 | Extended Data Fig. 4d |
| ySH31 | *MATa pCUP1-Ub-K-TRP1::NAT ubp6::KAN (pySH01: ubp6-I329A-L330A [CEN4/URA3])* | sJH138 | Extended Data Fig. 4d |
| ySH53 | *MATa pCUP1-Ub-K-TRP1::NAT ubp6::KAN (pySH07: ubp6-UBL [CEN4/URA3])* | sJH138 | Extended Data Fig. 4d |
| YTS119 | *MATa pCUP1-Ub-K-TRP1::HGR RPT1(WT)::NAT* | – | Extended Data Fig. 5f |
| YTS113 | *MATa pCUP1-Ub-K-TRP1::HGR RPT1(WT)::NAT ubp6::KAN* | – | Extended Data Fig. 5f |
| YTS159 | *MATa pCUP1-Ub-K-TRP1::HGR rpt1-S164R-T166K::NAT* | – | Extended Data Fig. 5f |
| YTS157 | *MATa pCUP1-Ub-K-TRP1::HGR rpt1-S164R-T166K::NAT ubp6::KAN* | – | Extended Data Fig. 5f |

**Supplementary Table 9.**
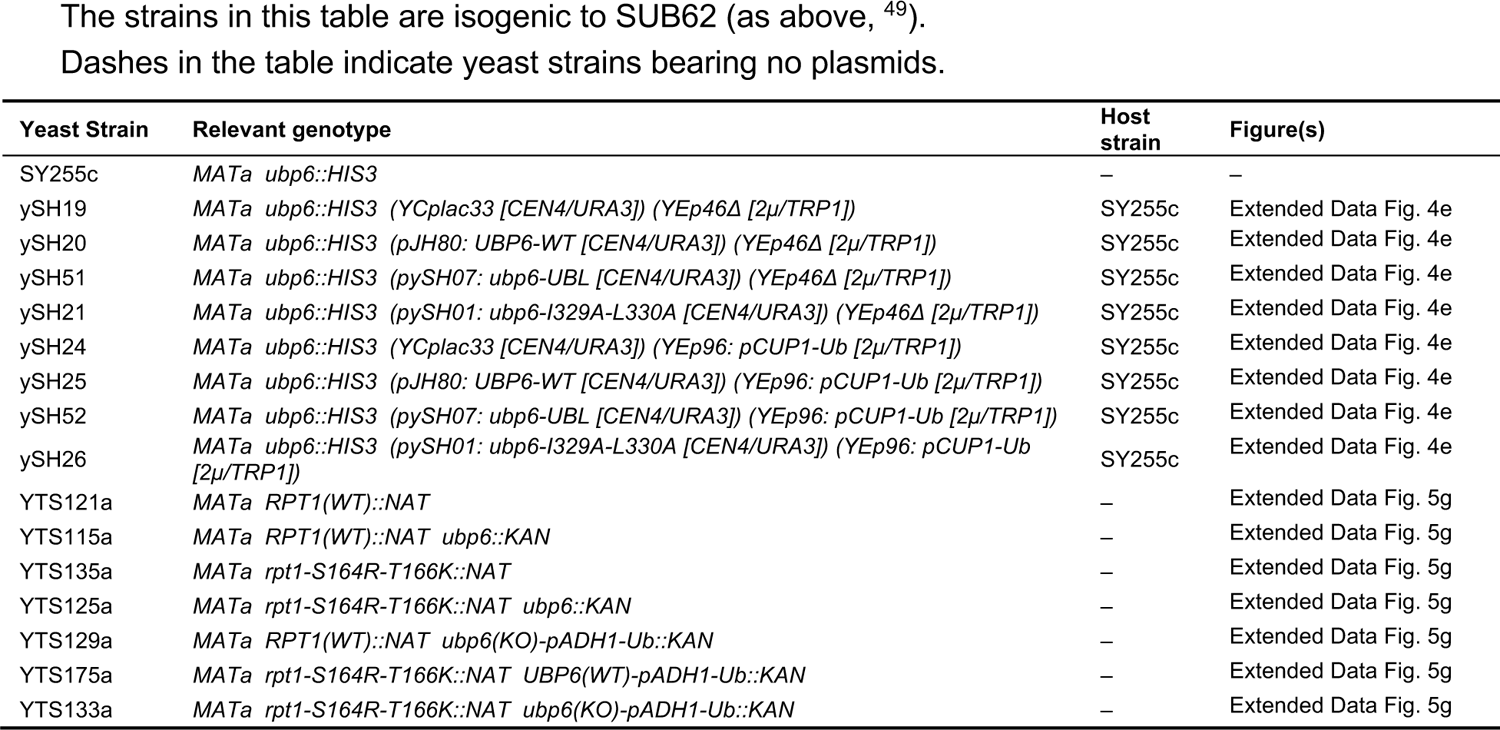
Yeast strains used for canavanine assay.

**Supplementary Table 10.** Yeast strains used for in vivo Ubp6 noncatalytic assay.

| Yeast Strain | Relevant genotype | Host strain | Figure(s) |
| --- | --- | --- | --- |
| sJH183<sup>48</sup> | *MATa ubp6::HIS3* | – | Extended Data Fig. 6a |
| ySH1 | *MATa ubp6::HIS3 (YCplac33 [CEN4/URA3])* | sJH183 | Extended Data Fig. 6a |
| ySH2 | *MATa ubp6::HIS3 (pJH80: UBP6-WT [CEN4/URA3])* | sJH183 | Extended Data Fig. 6a |
| ySH3 | *MATa ubp6::HIS3 (pySH01: ubp6-I329A-L330A [CEN4/URA3])* | sJH183 | Extended Data Fig. 6a |
| ySH6 | *MATa ubp6::HIS3 (pJH81: ubp6-C118A [CEN4/URA3])* | sJH183 | Extended Data Fig. 6a |
| ySH7 | *MATa ubp6::HIS3 (pySH04: ubp6-C118A-I329A-L330A [CEN4/URA3])* | sJH183 | Extended Data Fig. 6a |
| sJH185<sup>48</sup> | *MATa ubp6::HIS3 rpn4::KAN* | – | Extended Data Fig. 6a |
| ySH10 | *MATa ubp6::HIS3 rpn4::KAN (YCplac33 CEN4/URA3])* | sJH185 | Extended Data Fig. 6a |
| ySH11 | *MATa ubp6::HIS3 rpn4::KAN (pJH80: UBP6-WT [CEN4/URA3])* | sJH185 | Extended Data Fig. 6a |
| ySH12 | *MATa ubp6::HIS3 rpn4::KAN (pySH01: ubp6-I329A-L330A [CEN4/URA3])* | sJH185 | Extended Data Fig. 6a |
| ySH15 | *MATa ubp6::HIS3 rpn4::KAN (pJH81: ubp6-C118A [CEN4/URA3])* | sJH185 | Extended Data Fig. 6a |
| ySH16 | *MATa ubp6::HIS3 rpn4::KAN (pySH04: ubp6-C118A-I329A-L330A [CEN4/URA3])* | sJH185 | Extended Data Fig. 6a |
| SSW32 | *MATa ubp6::NAT rpn4::KAN* | – | Extended Data Fig. 6b |
| SSW272 | *MATa rpn4::KAN ubp6::NAT rpt1-S164R-T166K::NAT* | – | Extended Data Fig. 6b |

**Supplementary Table 11.** Reagents used in this study.

| Reagent | Source | Reference |
| --- | --- | --- |
| Rosetta 2 (DE3) competent cells | EMD Millipore | 71397-4 |
| IPTG | Gold Biotech | I2481C50 |
| AEBSF (200 mM stock, Water) | Gold Biotech | A5440 |
| Leupeptin | Sigma | L0649 |
| Antipain | Sigma | A6191 |
| Benzamidine hydrochloride | Sigma | 199001 |
| Aprotinin | Sigma | A6279 |
| Chymostatin | Sigma | C7268 |
| Pepstatin | Sigma | P5318 |
| Coomassie Plus Bradford Protein Assay | Pierce | PI23236 |
| Ni-NTA agarose | Qiagen | 30210 |
| Superdex 200 HiLoad 16/600 column | GE Healthcare | 28-9893-35 |
| Glutathione Sepharose 4B resin | GE Healthcare | 95017-172 |
| Thrombin protease | Sigma | T7513 |
| Benzamidine Sepharose | GE Healthcare | 17-0568-01 |
| BL21 (DE3) competent cells | EMD Millipore | 69450-3 |
| cOmplete, Mini, EDTA-free Protease Inhibitor Cocktail Tablets | Roche | 11836170001 |
| Amicon Ultra-4 Centrifugal Device, MWCO 30K | EMD Millipore | UFC803024 |
| Superdex 75 10/300 GL column | GE Healthcare | 17-5174-01 |
| ATP | Sigma | A3377 |
| M2 anti FLAG agarose | Sigma | A2220 |
| 3X FLAG peptide | Sigma | F4799 |
| Superose 6 10/300 GL column | GE Healthcare | 17517201 |
| Rabbit IgG resin | MP Biomedicals | 855961 |
| DTT | Gold Biotech | DTT50 |
| His-AcTEV protease (10000 U @ 10 U/uL) | Thermo Fisher | 12575-023 |
| NeutrAvidin agarose resin | Thermo Fisher | PI29201 |
| UbVS | Boston Biochem | U202 |
| Deuterium oxide D<sub>2</sub>O, low paramagnetic | Cambridge Isotope Laboratories | DLM-11-100 |
| ACQUITY UPLC BEH C18, 1.7 µm, 2.1 mm × 5 mm column | Waters | 186003975 |
| ACQUITY UPLC BEH C18, 1.8 µm, 1.0 mm × 100 mm column | Waters | 186002346 |
| Glu<sup>1</sup>-fibrinopeptide | Sigma | F3261 |
| ovalbumin | Sigma | A5503 |
| Low volume 384-well black flat bottom nonbinding surface microplate | Corning | 3820 |
| EnVision plate reader | Perkin Elmer | 2103 |
| Ub-AMC | Boston Biochem | U550 |
| suc-LLVY-AMC | Bachem | I-1395 |
| anti-Cdc27 (Clone AF3.1) | Santa Cruz | sc-9972 |
| Protein-A Sepharose 4B fast flow | Sigma | P9424 |
| Creatine kinase | Roche | 10127566001 |
| Creatine phosphate | Roche | 10621712001 |
| ADP-Potassium Salt<br>(0.25 M stock, 50 mM Tris-HCl [pH7.5], 5 mM MgCl<sub>2</sub>) | EMD Millipore | 117105 |
| Hexokinase (10 U/µL stock, water) | Sigma | H4502 |
| ATP<sub>γ</sub>S (10 mg/mL = 18.28 mM stock, water) | Santa Cruz | sc-214500A |
| PS-341/Veclade/Bortezomib (2.5 mM stock, DMSO) | ApexBio | A2614 |
| MG-262 (10 mg/mL = 20.4 mM stock, DMSO) | ApexBio | A8179 |
| Epoxomicin (10 mg/mL = 18.03 mM stock, DMSO) | ApexBio | A2606 |
| 1,10-phenanthroline (1.5 M stock, DMSO) | Sigma | 131377 |
| Lithium acetate | Sigma | L6883 |
| PEG MW 3350 | Sigma | P4338 |
| Salmon Sperm DNA (sheared, 10 mg/mL) | Agilent | 201190 |

## References

1. Groll, M. et al. A gated channel into the proteasome core particle. Nat Struct Biol 7, 1062–1067, doi:10.1038/80992 (2000).

2. Lenkinski, R. E., Chen, D. M., Glickson, J. D. & Goldstein, G. Nuclear magnetic resonance studies of the denaturation of ubiquitin. Biochim Biophys Acta 494, 126–130, doi:10.1016/0005-2795(77)90140-4 (1977).

3. Dong, Y. et al. Cryo-EM structures and dynamics of substrate-engaged human 26S proteasome. Nature 565, 49–55, doi:10.1038/s41586-018-0736-4 (2019).

4. Verma, R. et al. Role of Rpn11 metalloprotease in deubiquitination and degradation by the 26S proteasome. Science 298, 611–615, doi:10.1126/science.1075898 (2002).

5. Yao, T. & Cohen, R. E. A cryptic protease couples deubiquitination and degradation by the proteasome. Nature 419, 403–407, doi:10.1038/nature01071 (2002).

6. Bard, J. A. M., Bashore, C., Dong, K. C. & Martin, A. The 26S Proteasome Utilizes a Kinetic Gateway to Prioritize Substrate Degradation. Cell 177, 286–298 e215, doi:10.1016/j.cell.2019.02.031 (2019).

7. de Poot, S. A. H., Tian, G. & Finley, D. Meddling with Fate: The Proteasomal Deubiquitinating Enzymes. J Mol Biol 429, 3525–3545, doi:10.1016/j.jmb.2017.09.015 (2017).

8. Lee, B. H. et al. USP14 deubiquitinates proteasome-bound substrates that are ubiquitinated at multiple sites. Nature 532, 398–401, doi:10.1038/nature17433 (2016).

9. Aufderheide, A. et al. Structural characterization of the interaction of Ubp6 with the 26S proteasome. Proc Natl Acad Sci U S A 112, 8626–8631, doi:10.1073/pnas.1510449112 (2015).

10. Leggett, D. S. et al. Multiple associated proteins regulate proteasome structure and function. Mol Cell 10, 495–507 (2002).

11. Shi, Y. et al. Rpn1 provides adjacent receptor sites for substrate binding and deubiquitination by the proteasome. Science 351, doi:10.1126/science.aad9421 (2016).

12. Collins, G. A. & Goldberg, A. L. Proteins containing ubiquitin-like (Ubl) domains not only bind to 26S proteasomes but also induce their activation. Proc Natl Acad Sci U S A 117, 4664–4674, doi:10.1073/pnas.1915534117 (2020).

13. Hu, M. et al. Structure and mechanisms of the proteasome-associated deubiquitinating enzyme USP14. EMBO J 24, 3747–3756, doi:10.1038/sj.emboj.7600832 (2005).

14. Zhang, F. et al. Structural insights into the regulatory particle of the proteasome from Methanocaldococcus jannaschii. Mol Cell 34, 473–484, doi:10.1016/j.molcel.2009.04.021 (2009).

15. Hanna, J. et al. Deubiquitinating enzyme Ubp6 functions noncatalytically to delay proteasomal degradation. Cell 127, 99–111, doi:10.1016/j.cell.2006.07.038 (2006).

16. Bashore, C. et al. Ubp6 deubiquitinase controls conformational dynamics and substrate degradation of the 26S proteasome. Nat Struct Mol Biol 22, 712–719, doi:10.1038/nsmb.3075 (2015).

17. Eisele, M. R. et al. Expanded Coverage of the 26S Proteasome Conformational Landscape Reveals Mechanisms of Peptidase Gating. Cell Rep 24, 1301–1315 e1305, doi:10.1016/j.celrep.2018.07.004 (2018).

18. Wehmer, M. et al. Structural insights into the functional cycle of the ATPase module of the 26S proteasome. Proc Natl Acad Sci U S A 114, 1305–1310, doi:10.1073/pnas.1621129114 (2017).

19. Matyskiela, M. E., Lander, G. C. & Martin, A. Conformational switching of the 26S proteasome enables substrate degradation. Nat Struct Mol Biol 20, 781–788, doi:10.1038/nsmb.2616 (2013).

20. Clague, M. J., Urbe, S. & Komander, D. Breaking the chains: deubiquitylating enzyme specificity begets function. Nat Rev Mol Cell Biol 20, 338–352, doi:10.1038/s41580-019-0099-1 (2019).

21. Hu, M. et al. Crystal structure of a UBP-family deubiquitinating enzyme in isolation and in complex with ubiquitin aldehyde. Cell 111, 1041–1054, doi:10.1016/s0092-8674(02)01199-6 (2002).

22. Li, H. et al. Allosteric Activation of Ubiquitin-Specific Proteases by beta-Propeller Proteins UAF1 and WDR20. Mol Cell 63, 249–260, doi:10.1016/j.molcel.2016.05.031 (2016).

23. Samara, N. L. et al. Structural insights into the assembly and function of the SAGA deubiquitinating module. Science 328, 1025–1029, doi:10.1126/science.1190049 (2010).

24. Clerici, M., Luna-Vargas, M. P., Faesen, A. C. & Sixma, T. K. The DUSP-Ubl domain of USP4 enhances its catalytic efficiency by promoting ubiquitin exchange. Nat Commun 5, 5399, doi:10.1038/ncomms6399 (2014).

25. Avvakumov, G. V. et al. Amino-terminal dimerization, NRDP1-rhodanese interaction, and inhibited catalytic domain conformation of the ubiquitin-specific protease 8 (USP8). J Biol Chem 281, 38061–38070, doi:10.1074/jbc.M606704200 (2006).

26. Beckwith, R., Estrin, E., Worden, E. J. & Martin, A. Reconstitution of the 26S proteasome reveals functional asymmetries in its AAA+ unfoldase. Nat Struct Mol Biol 20, 1164–1172, doi:10.1038/nsmb.2659 (2013).

27. Leggett, D. S., Glickman, M. H. & Finley, D. Purification of proteasomes, proteasome subcomplexes, and proteasome-associated proteins from budding yeast. Methods Mol Biol 301, 57–70, doi:10.1385/1-59259-895-1:057 (2005).

28. Elsasser, S., Shi, Y. & Finley, D. Binding of ubiquitin conjugates to proteasomes as visualized with native gels. Methods Mol Biol 832, 403–422, doi:10.1007/978-1-61779-474-2_28 (2012).

29. Zheng, S. Q. et al. MotionCor2: anisotropic correction of beam-induced motion for improved cryo-electron microscopy. Nat Methods 14, 331–332, doi:10.1038/nmeth.4193 (2017).

30. Scheres, S. H. RELION: implementation of a Bayesian approach to cryo-EM structure determination. J Struct Biol 180, 519–530, doi:10.1016/j.jsb.2012.09.006 (2012).

31. Rohou, A. & Grigorieff, N. CTFFIND4: Fast and accurate defocus estimation from electron micrographs. J Struct Biol 192, 216–221, doi:10.1016/j.jsb.2015.08.008 (2015).

32. Wagner, T. et al. SPHIRE-crYOLO is a fast and accurate fully automated particle picker for cryo-EM. Commun Biol 2, 218, doi:10.1038/s42003-019-0437-z (2019).

33. Adams, P. D. et al. PHENIX: building new software for automated crystallographic structure determination. Acta Crystallogr D Biol Crystallogr 58, 1948–1954, doi:10.1107/s0907444902016657 (2002).

34. Leaver-Fay, A. et al. ROSETTA3: an object-oriented software suite for the simulation and design of macromolecules. Methods Enzymol 487, 545–574, doi:10.1016/B978-0-12-381270-4.00019-6 (2011).

35. Humphrey, W., Dalke, A. & Schulten, K. VMD: visual molecular dynamics. J Mol Graph 14, 33–38, 27-38, doi:10.1016/0263-7855(96)00018-5 (1996).

36. Schweitzer, A. et al. Structure of the human 26S proteasome at a resolution of 3.9 A. Proc Natl Acad Sci U S A 113, 7816–7821, doi:10.1073/pnas.1608050113 (2016).

37. Pettersen, E. F. et al. UCSF Chimera--a visualization system for exploratory research and analysis. J Comput Chem 25, 1605–1612, doi:10.1002/jcc.20084 (2004).

38. Goh, B. C. et al. Computational Methodologies for Real-Space Structural Refinement of Large Macromolecular Complexes. Annu Rev Biophys 45, 253–278, doi:10.1146/annurev-biophys-062215-011113 (2016).

39. Phillips, J. C. et al. Scalable molecular dynamics with NAMD. J Comput Chem 26, 1781–1802, doi:10.1002/jcc.20289 (2005).

40. Ribeiro, J. V. et al. QwikMD - Integrative Molecular Dynamics Toolkit for Novices and Experts. Sci Rep 6, 26536, doi:10.1038/srep26536 (2016).

41. Word, J. M., Lovell, S. C., Richardson, J. S. & Richardson, D. C. Asparagine and glutamine: using hydrogen atom contacts in the choice of side-chain amide orientation. J Mol Biol 285, 1735–1747, doi:10.1006/jmbi.1998.2401 (1999).

42. Liebschner, D. et al. Macromolecular structure determination using X-rays, neutrons and electrons: recent developments in Phenix. Acta Crystallogr D Struct Biol 75, 861–877, doi:10.1107/S2059798319011471 (2019).

43. Emsley, P., Lohkamp, B., Scott, W. G. & Cowtan, K. Features and development of Coot. Acta Crystallogr D Biol Crystallogr 66, 486–501, doi:10.1107/S0907444910007493 (2010).

44. Scheurer, M. et al. PyContact: Rapid, Customizable, and Visual Analysis of Noncovalent Interactions in MD Simulations. Biophys J 114, 577–583, doi:10.1016/j.bpj.2017.12.003 (2018).

45. Williams, C. J. et al. MolProbity: More and better reference data for improved all-atom structure validation. Protein Sci 27, 293–315, doi:10.1002/pro.3330 (2018).

46. Dimova, N. V. et al. APC/C-mediated multiple monoubiquitylation provides an alternative degradation signal for cyclin B1. Nat Cell Biol 14, 168–176, doi:10.1038/ncb2425 (2012).

47. Weissmann, F. et al. biGBac enables rapid gene assembly for the expression of large multisubunit protein complexes. Proc Natl Acad Sci U S A 113, E2564–2569, doi:10.1073/pnas.1604935113 (2016).

48. Hanna, J., Meides, A., Zhang, D. P. & Finley, D. A ubiquitin stress response induces altered proteasome composition. Cell 129, 747–759, doi:10.1016/j.cell.2007.03.042 (2007).

49. Finley, D., Ozkaynak, E. & Varshavsky, A. The yeast polyubiquitin gene is essential for resistance to high temperatures, starvation, and other stresses. Cell 48, 1035–1046, doi:10.1016/0092-8674(87)90711-2 (1987).

50. Saeki, Y. et al. Knocking out ubiquitin proteasome system function in vivo and in vitro with genetically encodable tandem ubiquitin. Methods Enzymol 399, 64–74, doi:10.1016/S0076-6879(05)99005-8 (2005).

51. Gietz, R. D. & Sugino, A. New yeast-Escherichia coli shuttle vectors constructed with in vitro mutagenized yeast genes lacking six-base pair restriction sites. Gene 74, 527–534 (1988).

52. Ecker, D. J., Khan, M. I., Marsh, J., Butt, T. R. & Crooke, S. T. Chemical synthesis and expression of a cassette adapted ubiquitin gene. J Biol Chem 262, 3524–3527 (1987).

53. Brachmann, C. B. et al. Designer deletion strains derived from Saccharomyces cerevisiae S288C: a useful set of strains and plasmids for PCR-mediated gene disruption and other applications. Yeast 14, 115–132, doi:10.1002/(SICI)1097-0061(19980130)14:2<115::AID-YEA204>3.0.CO;2-2 (1998).

